# *Acropora millepora*’s microbiome is predicted by algal symbionts, host genetics, and environment

**DOI:** 10.64898/2025.12.15.694400

**Authors:** Karim Primov, Carly Scott, Alexa Huzar, Christopher Peterson, Mark Kirkpatrick, Mikhail Matz

## Abstract

The coral microbiome is a critical component of coral health and resilience, yet it is unclear what factors drive coral microbiome composition, especially within the context of coral bleaching. Here, we use whole genome sequencing data combined with a machine learning approach (RDAforest) to assess predictors of the microbiome in 208 colonies of *Acropora millepora* from 12 reef sites in the Central Great Barrier Reef during a 2016 bleaching event. We characterized microbiome variation using k-mers. While some environmental variables, such as chlorophyll seasonal range and maximum degree heating weeks, were associated with microbiome composition, we find that host genetics and dominant photosymbionts were more powerful predictors. In contrast, bleaching score had negligible predictive power. The coral’s microbiome therefore correlates with dominant photosymbiont identity even during a bleaching event. The association of the microbiome with the environment suggests that the coral microbiome can serve as a proxy for environmental variation when environment cannot be measured directly, which may be especially useful in ancient DNA studies.

## 1. Introduction

Various stressors related to climate change have led to coral decline worldwide (Donner *et al*., 2005; De’ath *et al*., 2012; Jones *et al*., 2008; Glynn *et al*., 1996; McClanahan 2004; Sully *et al*., 2019; West & Salm 2003). The Great Barrier Reef, the largest barrier reef system on the planet, has been undergoing severe decline in coral cover and diversity in recent decades (Ceccarelli *et al*., 2024; De’ath *et al*., 2012; Emslie *et al*., 2024). Coral bleaching, the dysbiosis between the coral host and its photosymbiotic zooxanthellae largely driven by heat stress, varies across biogeographic regions, coral taxa, and environments (Marshall *et al*., 2000; Baker *et al*., 2008; Barshis *et al*., 2013; Swain *et al*., 2016; Hughes *et al*., 2018; Sully *et al*., 2019; van Woesik *et al*., 2022). The majority of coral adaptation and bleaching variation research has focused on coral host and/or symbiont genetic background (Black *et al*., 2025; Cabacungan *et al*., 2025; Fuller *et al*., 2020, Gallery *et al*., 2025; Rippe *et al*., 2021; Scott *et al*., 2025; Sturm *et al*., 2020, 2022, 2022). Few studies consider the bacterial microbiome as a potential driver of bleaching variation. Previous work has focused on genomic predictions of bleaching responses in central

Great Barrier Reef *Acropora millepora* which exhibit inter- and intrapopulation bleaching variation (Figure 1; Fuller *et al*., 2020). Little host population structure was detected, while colonies with higher *Durusdinium* photosymbiont proportions bleached less than those dominated by *Cladocopium*. Bleaching response was found to be polygenic, and Fuller *et al*. (2020) concluded that integrating host and symbiont genetic data, as well as environmental data, can aid in distinguishing colonies that are more tolerant to bleaching than others. The coral microbiome’s role in bleaching and symbiont community variation still remains largely unknown.

**Figure 1:**
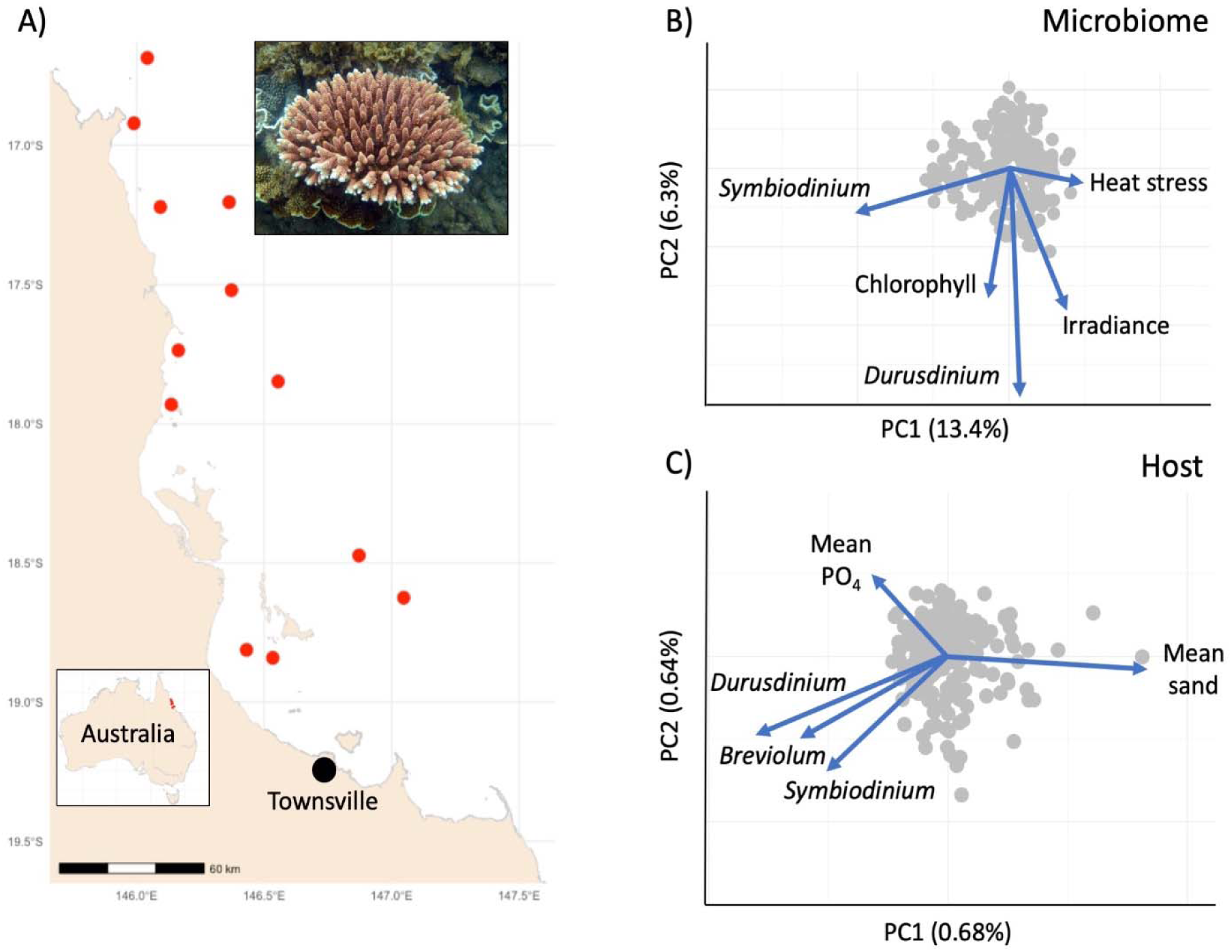
A: Sampling sites of *Acropora millepora* from Fuller *et al*. (2020). B: Top five predictors of microbiome variation aligned to its unconstrained ordination based on euclidean distances (after centered-log-ratio transformation). C: Top five predictors of host genetic variation aligned to its unconstrained ordination based on identity-by-state distances. For both (B) and (C), the predictors were identified by RDAforest analysis.

The coral bacterial microbiome has recently been proposed to be crucial for coral host development, survival, and climate stress resilience (Delgadillo-Ordonez *et al*., 2024; Peixoto *et al*., 2017; Rosado *et al*., 2019; Santoro *et al*., 2021; Voolstra *et al*., 2024). The coral microbiome has been found to be influenced by the coral host, the environment, as well as bleaching severity (Gardner *et al*., 2019; Scott *et al*., 2025). Coral microbiomes are also intricately associated with corals’ algal photosymbionts, as intracellular bacteria have been found inside zooxanthellae (Maire *et al*., 2021). It is still unresolved, however, whether the coral microbiome plays a role in determining bleaching susceptibility or zooxanthellae community assembly. Microbiome shifts during bleaching events are well documented, however it is unclear whether the shifts are causal, correlational, or a direct consequence of bleaching. Disentangling the major drivers of microbiome variation from factors such as underlying host genetic background has been largely understudied and is a necessary step to adequately assess the relative contribution of the microbiome to bleaching & symbiont community variation. This is of utmost importance in reef systems such as the Florida Keys and the Great Barrier Reef which have both undergone unprecedented declines in both coral diversity and cover in recent decades.

Beyond the role the microbiome may play in bleaching, there is also increased interest in using the coral microbiomes to characterize environmental conditions in threatened coral reefs. Traditionally, restoration efforts in the threatened coral reef habitats have used satellite-derived measurements of environmental variables to characterize coral reef environmental conditions. There has been a shift, however, to assessing the diagnostic capacity of the microbiome of various reef-dwelling taxa, and the surrounding seawater, to assess environmental conditions and incipient change of environmental conditions on a coral reef (Glasl *et al*., 2019). Previous efforts to assess the diagnostic capacity of coral and seawater microbiomes did not control for underlying host genetic background and symbiont community variation, which are two factors known to predict microbiome variation.

In the current study, we leverage the breadth and depth of whole genome sequencing data, bleaching data, symbiont proportions, and environmental data from Fuller *et al*., (2020) to assess various aspects of *Acropora millepora*’s microbiome and host genetics with respect to bleaching and symbiont community variation. We test whether the microbiome is primarily influenced by bleaching status or by the environment, host genetics, and dominant photosymbiont identity. While the vast majority of published coral microbiome studies relied on 16S amplicon sequencing, here we detect microbial taxa based on k-mer occurrence in whole-genome coral sequencing reads, an approach that is expected to be both more comprehensive and less biased. We then use a random forest based approach (Breiman 2001) to identify the most prominent predictors of microbiome composition. We identify environmental variables associated with overall host genetic variation, and we assess the possibility of using the microbiome to infer biologically relevant environmental variables.

## 2. Methods

### 2.1 Data curation, microbial community characterization, and multilocus genotyping

Fuller *et al*. (2020) sampled 237 colonies of *Acropora millepora* across 12 populations throughout the Central Great Barrier Reef during the 2016/2017 bleaching event (Figure 1A). Raw reads for these samples were retrieved from NCBI (Bioproject PRJNA593014). After removal of outliers samples that were likely species misidentifications and samples with high relatedness (likely clones), 208 whole genome sequence samples with coverage ranging from 1x to 7x were used for bacterial microbial ecology analysis. Reads were trimmed and filtered to remove smaller and low quality reads using TrimGalore (Krueger 2015), then aligned to the *Acropora millepora* reference genome (Fuller *et al*., 2020) to remove coral-aligned reads. Unaligned reads were then aligned to updated reference genomes for the four major coral photosymbiont genera (*Symbiodinium microadriaticum* [http://smic.reefgenomics.org/download/], *Breviolum minutum* [https://www.ncbi.nlm.nih.gov/datasets/genome/GCA_000507305.1/], *Cladocopium goreaui* [https://www.ncbi.nlm.nih.gov/datasets/genome/GCA_947184155.2/]*, and Durusdinium trenchii* [https://www.ncbi.nlm.nih.gov/datasets/genome/GCA_963970005.1/]) to assess dominant photosymbiont composition and to remove symbiont reads from the data. Filtered reads were then taxonomically profiled using Kraken2, with the Standard-16 reference database (Wood *et al*., 2019). The standard-16 database contains archaeal, bacterial, viral, plasmid, human, and Univec_core (common primers, adapters, linkers, and vectors) reference sequences, capped at 16 Gb to limit computational usage. Bracken (Lu *et al*., 2017) was used to re-estimate Kraken2 family-level bacterial abundances.

Coral-aligned reads were used to generate the host genetic identity-by-state (IBS) distance matrix using ANGSD (Korneliussen *et al*. 2014), using loci genotyped in at least 50% of samples.

Family-level microbiome profiles were filtered to remove eukaryotic, viral, plasmid, and archaeal taxa. The remaining bacterial abundances were centered-log-ratio transformed and used to generate a distance matrix based on Euclidean distances. This distance matrix was the basis for the ordination used in RDAforest analysis.

Raw reads were also aligned to an *Endozoicomonas acroporae* genome (Tandon *et al*., 2020) concatenated to *Acropora millepora*’s chromosome 14 (the smallest chromosome) to calculate the relative abundance of *Endozoicomonas*.

### 2.2 RDAforest

RDAforest (Matz and Black 2024) was used to assess the most significant predictors of 1) microbiome variation, 2) host genetic variation, 3) variation in the relative abundance of the putative beneficial bacterium *Endozoicomonas*, and 4) the microbial families most predictive of environmental variation. RDAforest relies on random forest regression to 1) identify the importance, or predictive power, of each predictor variable while accounting for all possible interactions between predictors, **2)** identify the most likely primary driver among a group of several correlated predictors, and **3)** assess the uncertainty in estimating the importances of individual predictors.

When identifying the major predictors of microbiome variation, we wanted to also measure the drop in model performance when each of the explanatory metadata categories (bleaching scores, environmental data, first 20 host genetic principal components, population/site identity, symbiont community proportions; Figure 2c) was withheld from the analysis. For bleaching scores, samples with missing bleaching scores (43/208; 21%) were imputed either with the average bleaching score or using the program *missForest* (Stekhoven & Bühlmann 2012).

**Figure 2:**
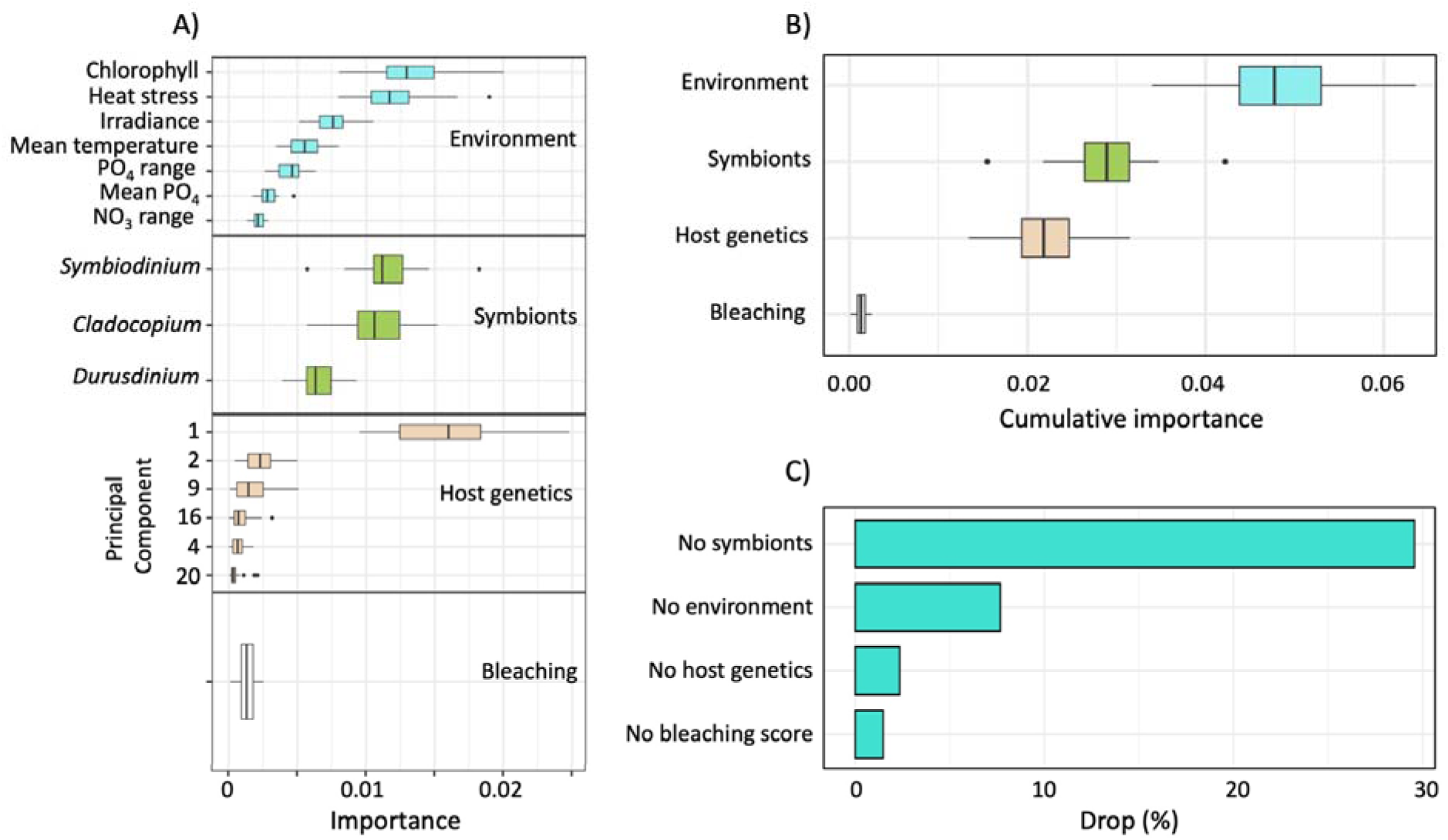
A: Strongest predictors of microbiome variation grouped by metadata category. B: Cumulative importances of metadata categories explaining microbiome variation in the full RDAForest model. C: Drop in model performance when omitting each metadata category from the model.

### 2.3 Microbial predictors of environmental variation

RDAforest was again used to measure the percent of environmental variation predicted by the overall microbiome. To assess the association of specific microbial taxa, we performed linear regression of their centered-log-ratio transformed abundances on the first two principal components of environmental variation across 12 reef sites. For this analysis, highly correlated environmental variables were identified using the function “hclust” in the R package “stats”, and only variables that had Pearson correlation *r* < 0.8 to each other were selected (Figure S1), choosing one representative variable for groups of variables with *r* ≥ 0.8. The same procedure was performed for selected microbial taxa that correlate with PC1 and PC2, to assess whether these selected microbial families are correlated with each other (Figure S2). A PCA ordination of sampled sites was constructed based on retained environmental variables. Indicator microbial taxa were then identified as highly significantly associated (by linear regression, *p* < 0.001) with any of the first two environmental PCs, using the function “envfit” in the R package “vegan”. The first two PCs together explained 63% of total environmental variation.

### 2.4 Correlations and non-parametric regressions

To assess linear relationships between various measures and predictors, the function “cor.test” in the “stats” package in R was used. Non-parametric regressions were performed using the function “gam” in the “mgcv” package in R. The function “protest” of the “vegan” package in R was also used to assess the multivariate similarity between host genetic variation and bacterial microbiome variation.

## 3. Results

### 3.1 Dominant photosymbionts and microbes

After realignment of raw reads to updated reference genomes for the four major photosymbiont genera (*Symbiodinium microadriaticum*, *Breviolum minutum*, *Cladocopium goreaui*, and *Durusdinium trenchii*), *Cladocopium* and *Durusdinium* were identified as the overwhelmingly dominant genera across all samples, with 91.3% (190/208) of samples having either genus represent >90% of their photosymbiont profiles. *Cladocopium* and *Durusdinium*-dominated samples harbored smaller proportions (<16%) of the other symbiont genera. *Symbiodinium* was a minor symbiont, with only 12/208 samples containing >1% *Symbiodinium* and only 4/208 samples containing >10% *Symbiodinium* photosymbionts. Meanwhile, *Breviolum* was nearly absent from all samples, with only 1/208 samples containing >1% *Breviolum* and no samples containing >10% *Breviolum* photosymbionts.

After microbial taxonomic profiling using Kraken2 and Bracken, the dominant microbial families include *Streptomycetaceae*, *Pseudomonadaceae*, *Enterobacteriaceae*, *Bacillaceae*, and Flavobacteriaceae (Figure S3). *Endozoicomonas,* within the family *Endozoicomonadaceae*, was among the top 25 most abundant microbial families (Figure S3).

### 3.2 Predictors of the microbiome

The proportion of total microbiome variation explained by all predictor variables by RDAForest’s gradient forest analysis was 16%. After RDAforest’s predictor selection procedure (function “mtrySelJack”) which is designed to discard collinear predictors, 17 predictors were retained out of the original 70, together explaining 11% of total microbiome variation (Figure 2A, 2B, S5). The top five strongest factors predictors were host genetic PC1, chlorophyll-a seasonal range, maximum degree heating weeks, *Symbiodinium* proportions, and *Durusdinium* proportions (Fig. 1B, 2A, 2B, S5). Collectively, environmental variables explained the largest proportion of microbiome variation due to the large number of environmental variables, followed by symbiont proportions, host genetics, and bleaching score (Figure 2B).

RDAForest’s turnover curves for individual predictors reveal where, along the range of predictor values, changes in the microbiome occur. A notable transition occurred at ∼0.5 mg/m^3^ of chlorophyll-a seasonal range, while a less pronounced transition was observed at ∼7.6 degree heating weeks (Figure S6). Transitions were observed as *Cladocopium* changed in relative proportion starting at 20% and for *Durusdinium* at ∼10%, and as *Symbiodinium* increased in relative proportion starting at 0.75% (Figure S6).

Runs of RDAForest that excluded individual metadata categories revealed that removing symbiont community proportions resulted in the largest predictive power drop (∼30% of full model’s power, Figure 2C, S7; Table S2). This indicates that symbiont communities are the strongest predictor of microbiome variation among the tested metadata categories. Dropping environmental metadata led to the second largest drop in model performance (7.7% of full model power), followed by host PCs (2.3%) and bleaching score (1.5%; Figure 2C, S7).

Procrustes test (function vegan::protest) revealed a highly significant similarity between the microbiome distance matrix and the host genetic IBS matrix (r = 0.45, *p* = 0.001). Both Pearson and Spearman-rho correlation tests revealed a significant correlation between the first principal component of the host genetics and the first principal component of microbiome variation (r = 0.61, p-value = 2.2 e-16; rho = 0.6811688, p-value = 2.2 e-16; Table S1). These results indicate that microbiome composition is structured by host genetic variation.

### 3.3 Host genetic predictors

The original analysis of Fuller *et al*. (2020) followed the GWAS paradigm: it relied on imputed coral genotypes and looked for individual SNP associations with bleaching and environmental predictors. Here, we reanalyzed the same data using RDAforest, which aims to detect highly polygenic responses, possibly undetectable at single-SNP level, by associating the overall genetic structure with predictors (Matz and Black 2024). Instead of imputed genotypes we used an entirely model-free genotyping approach (identity-by-state based on single read resampling, Korneliussen *et al*. 2014) for this analysis.

Our analysis revealed several variables together explain 38% of variation along the host PC1: *Symbiodinium* proportion, mean sand cover, *Durusdinium* and *Breviolum* photosymbiont proportions, and phosphate seasonal range (Figure 1C, S8). Host PC1 explains only 0.7% of total genetic variation. Host genetic variation is largely decoupled from the environment, bleaching score, and dominant photosymbiont identity, which is consistent with the results of Fuller *et al*., (2020).

### 3.4 The microbiome predicts environmental variables

RDAforest’s gradient forest analysis revealed that the top five strongest microbial predictors of environmental variation are Alcaligenaceae, Endozoicomonadaceae, Haliscomenobacteraceae, Erwiniaceae, and Enterobacteriaceae (Figure 3A). Collectively, the entire *A. millepora* microbiome explains >30% of environmental variation (Figure 3A).

**Figure 3:**
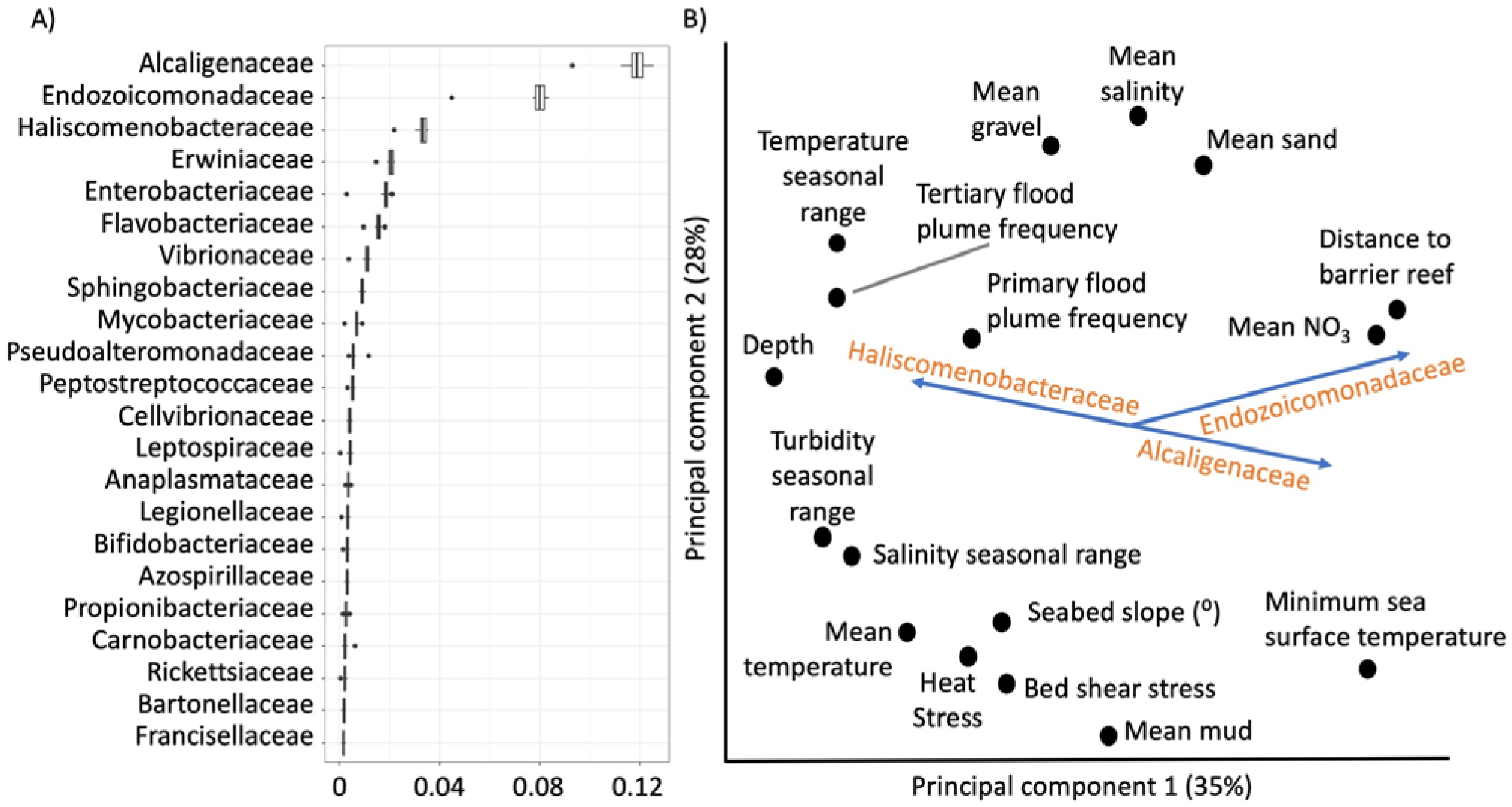
A: Microbial families most predictive of total environmental variation, according to RDAforest analysis. B: Microbial predictors of specific environmental variables. Points indicate scores of individual environmental predictors in unconstrained ordination of environmental variation. The arrows show alignment of abundances of top three microbial families that were predictive of the environment.

The envfit ordination (PC1: 35%, PC2: 28%; Figure 3B) revealed that environmental variables can be predicted by the abundance of specific microbial families. Endozoicomonadaceae is positively associated with distance to barrier reef and average nitrate concentrations and negatively associated with degree heating weeks (Figure 3B). Haliscomenobacteraceae is positively correlated with depth, while Alcaligenaceae is negatively correlated with depth (Figure 3B).

### 3.5 *Endozoicomonas* is found at low abundances and predicted by the environment

*Endozoicomonas*, the putatively beneficial coral microbe, was found at relatively low abundances (ranging from 0 to 105,247 reads per sample). The proportion of total *Endozoicomonas* variation explained by all predictor variables in RDAforest analysis was 63%. The top five predictors, explaining 25% of total variation, were benthic irradiance, average phosphate and temperature, and phosphate and nitrate seasonal ranges (Figure S10.). Overall, 14 out of 70 initial variables were chosen by RDAforest as significant predictors of *Endozoicomonas* abundance, collectively predicting ∼28% of its variation (Figure S10).

Turnover curves show changes in *Endozoicomonas* abundances for average temperature at and above 25.4 C, benthic irradiance at 0.09 mol photons m ² day ¹, average phosphate of 0.158 µM, average nitrate at 0.235 µM, phosphate seasonal range of 0.091 µM, and nitrate seasonal range of 0.245 µM (Figure S11). All other variables explained minimal *Endozoicomonas* variation.

### 3.6 Bleaching is a significant but weak predictor of microbiome and host genetic variation

Bleaching, despite explaining very little microbiome variation (Fig. 2A, 2B), was correlated with *Breviolum, Cladocopium*, *Durusdinium*, and *Symbiodinium* proportions, degree heating weeks, and *Endozoicomonas* proportions (Pearson correlation for all, *p* < 0.05; Tables S1). In contrast, Spearman-rho correlation tests failed to detect significant relationships between bleaching and both *Cladocopium* and *Durusdinium* proportions (Table S1). Non-parametric regression further indicated a significant relationship between bleaching and both *Cladocopium* and *Durusdinium* proportions (*p* < 10^-4^; Table S3).

Forty three samples with missing bleaching scores were imputed either with the average bleaching score, or using the program *missForest*, which produced very similar RDAforest results.

## 4. Discussion

*Acropora millepora*’s microbiome was best predicted by Symbiodinaceae community composition, followed by environmental variables and host genetic background, and (to a small extent) by bleaching status. The putatively beneficial bacteria genus, *Endozoicomonas*, was found at low relative abundances, and Endozoicomonadaceae abundance was among the best predictors of environmental variation. Overall, almost 30% of environmental variation can be predicted by microbiome composition.

### 4.1 Link between Symbiodiniaceae and microbiome

Symbiodiniaceae community composition and environmental variables were similarly predictive of coral microbiome variation according to the RDAforest procedure, which is based on randomizing each predictor individually (Fig. 2 A&B). Testing the importance of whole categories of predictors by omitting them from the model revealed that Symbiodiniaceae together provided the most explanatory power (Fig. 2C). Coral microbiomes have been previously found to differ depending on dominant photosymbiont identity. *Pocillopora acuta* off Hainan Island which were either *Durusdinium* or *Cladocopium* dominant were found to harbor distinct microbiomes. Microbiomes varied with respect to symbiont profiles in *Diploastrea heliopora*, *Pachyseris speciosa*, *Pocillopora acuta*, and *Porites lutea* in Singapore (Yang *et al*., 2025; Rabbani *et al*., 2025).

Microbiome variation has previously been found to be shaped by primary symbiosis, or ruderal microbial or eukaryotic symbiont occupancy, in other taxa including the bean bug *Riptortus pedestris,* the squash bug *Anasa tristis*, the barrelclover *Medicago trunculata,* and the fungus *Diploschistes lichenicolous* (Shan *et al*., 2024; Chen *et al*., 2024; Wang *et al*., 2021; Wedin *et al*., 2016). In addition to the photosynthate used by the coral host, Symbiodiniaceae produce amino acids, pigments, carotenoids, and steroid precursors such as squalenes and lanosterols (Mohamed *et al*., 2023). These exudates, the composition of which likely varies depending on Symbiodinaceae community, would influence trophic niches for the microbiome, leading to the observed link between the two.

The association between coral-hosted Symbiodiniaceae and microbes is well documented. In some cases, bacteria are present even inside zooxanthellae, and these intracellular bacteria are taxonomically diverse yet highly conserved across photosymbionts (Maire *et al*., 2021). Microbes positively impact Symbiodinaceae core photosynthetic health and efficiency, with some thermotolerant zooxanthellae harboring more stable microbiomes than thermosensitive zooxanthellae (Diaz-Almeyda *et al*., 2022; Matthews *et al*., 2023). Symbiodiniaceae community composition may exert an even greater influence on microbiome composition than environmental differences, as is the case for the sea anemone *Anthopleura elegantissima* (Morelan *et al*., 2019). In a review of bacterial-Symbiodiniaceae interactions, Symbiodiniaceae can be difficult to culture axenically, and Symbiodiniaceae genomes are relatively small compared to other dinoflagellates, suggesting the potential loss of production of key enzymes due to interactions with the coral host, microbiome, or both (Matthews *et al*., 2020). Coral photosymbionts, therefore, may have an intricate evolutionary relationship with microbes that could provision their photosymbionts with certain traits or functions that they would not have otherwise.

It remains unclear to what extent microbiome-shaping variation may be present within both *Cladocopium* and *Durusdinium*. The genus *Cladocopium* is notably diverse, likely containing hundreds of species, while *Durusdinium* also contains multiple species. Future work should employ higher-resolution Symbiodiniaceae genetic markers, such as *psb*A minicircle non-coding region (*psb*A), microsatellites (Butler *et al*., 2023), or, ultimately, SNPs, to answer this question.

### 4.2 Host genetics predicts microbiome

Interspecific differences in coral microbiomes are well documented. *Porites lobata*, *Pachyseris speciosa*, and *Merulina ampliata* consistently harbored distinct microbiomes from one another at both Kusu and Sisters Reefs in Singapore (Quek *et al*., 2023). *Acropora hemprichii* and *Pocillopora verrucosa* transplanted across sites varying in anthropogenic impact exhibited significantly distinct microbiomes from one another (Ziegler *et al*., 2019). *Acropora prostrata*, *Montipora digitata*, *Pocillopora acuta*, and *Porites cylindrica* on the same reef in Fiji also harbored distinct microbiomes (Longley *et al*., 2024). *Acropora millepora* and *A. tenuis* were also found to harbor host-specific microbiomes in the Great Barrier Reef (Glasl *et al*., 2019). To the best of our knowledge, however, there are no reports of intraspecific genetic variation influencing microbiomes in corals. Here, we found that host genetic PC1 was the strongest individual predictor of microbiome composition (Fig. 2 A), even despite minimal host population structure (the host genetic PC1 explains only 0.68% of total genetic variation). Another demonstration of the link between host genetic structure and microbiome composition is a highly significant Procrustes correlation (measure of concordance between two multivariate datasets) between them. The finding that even very subtle intraspecific genetic structure of the host is associated with shifts in microbiome composition is very intriguing, since microbiome can modify coral fitness (Voolstra et al., 2024) and thus influences the strength and/or direction of natural selection as populations diverge.

### 4.3 Environment and coral microbiome

The microbiome shifts when chlorophyll-a seasonal range is at and above ∼0.5 mg/m^3^ (Figure S6). While the dependence between microbiome and specific chlorophyll concentration is unclear, many studies find that cnidarian microbiomes shift in response to changes in productivity and seasonality, as is the case for the jewel sea anemone *Corynactis viridis* and the coral *Pavona decussata* (Palladino *et al*., 2021; Xu *et al*., 2024). Given the sensitivity of microbiome to environmental fluctuations (Glasl *et al*., 2019), the coral microbiome may be a useful diagnostic tool for assessing environmental fluctuations.

Degree heating weeks (DHW) showed two thresholds where microbiome composition changed: smaller change at ∼7.6 DHW and a larger change at ∼9.2 DHW (Figure S6). Reefs experiencing ≥ 8 DHW tend to show severe bleaching (Kayanne 2017; Skirving *et al*., 2019). Other studies showed that the microbiome shifts under thermal stress in *Porites lutea* from both the Andaman Sea and Pelorus Island off the Great Barrier Reef (Pootakham *et al*., 2019; Marangon *et al*., 2025). However, heat stress, in combination with acidification stress, did not significantly alter the microbiome of *Acropora millepora* from Fiji (Grotolli *et al*., 2018). These differences in microbiome response could be due to species-specific microbial responses, or differences in sampling size, or the extent of heat stress. Future research should investigate microbiome shifts in response to heat stress utilizing a standardized approach across multiple coral species.

### 4.4 Bleaching is a minor predictor of microbiome variation

Bleaching status was found to be the least influential predictor of microbiome variation, contrary to our initial hypothesis (Figures 2 & S5). Not all our samples had bleaching scores (165 of 208 samples), and so the missing scores had to be imputed, which may partially account for the lack of predictive power. Still, the result did not depend on the method of imputation (average bleaching score or *missForest* imputations).

Other studies have reported conflicting evidence about the role of bleaching in predicting the microbiome. *Porites lutea* microbiome variation in Andaman Sea throughout the 2016 bleaching event depended mainly on reef site and not on bleaching status. This may be influenced by cryptic genetic lineages partitioned by habitat within *Porites lutea*, with cryptic lineages being a common trend found within massive *Porites* and many corals (Black *et al*., 2025; Cabacungan *et al*., 2025; Primov *et al*., 2024; Rippe et al., 2021; Rivera *et al*., 2022; Scott *et al*., 2025; Starko *et al*., 2020). In another study, the microbiomes of *Acropora digitifera*, *Galaxea fascicularis* and *Porites pukoensis* in Hainan Island varied primarily by species identity, not bleaching condition (Xu *et al*., 2023). Bleaching, however, did cause significant changes in the microbiome across 53 species spanning 26 coral genera during a 2019 bleaching event in the Nansha Islands in the South China Sea (Sun *et al*., 2022). Future studies should assess the changes in the microbiome throughout the various stages of bleaching while controlling for host genetic background and assess the impact of any changes in the microbiome to changes in host fitness to properly disentangle the effects of the microbiome on bleaching status and coral health.

### 4.5 The microbiome is predictive of environmental variation

While the environment predicts only about 5% of microbiome variation, the reverse model, predicting the environment using microbiome, has much stronger predictive power: microbiome explains about 30% of environmental variation. This asymmetry is not surprising: many microbes are not associated with the environment and so are not predictable by it, whereas the reverse task relies only on strongly environment-associated microbes (Fig. 3 A&B).

Alcaligenaceae, the microbial family that best predicts environmental variation (12%, Fig. 3 A), correlates negatively with depth (Figure 3 B). Few studies report Alcaligenaceae as a coral-associated microbe, but it has been found at low pH in *Stylophora pistillata* and has also been identified as a denitrifying bacteria (Muniz-Barreto *et al*., 2021). The putatively beneficial microbe *Endozoicomonas*, despite being found at relatively low abundances, was the second-best predictor of environmental variation in RDAforest analysis (Fig. 3A). *Endozoicomonas* was found to positively correlate with average nitrate concentrations. This result is consistent with previous research finding that *Endozoicomonas* species including *E. numazuensis* and *E. promiscua* have been found to reduce nitrate to ammonia (Modolon *et al*., 2025; Nishijima *et al*., 2013; da Silva *et al*., 2025). *Endozoicomonas* abundance was also negatively correlated with maximum degree heating weeks. This result is consistent with previous findings of *Endozoicomonas*’ decreased abundance during bleaching and stressful climatic events, including *Acropora millepora* and *Porites cylindrica* found at carbon seeps in Papua New Guinea (Morrow *et al*., 2015) and *Pocillopora acuta* during a 2016 Great Barrier Reef bleaching event (Botte *et al*., 2022). Under non-stressful conditions, however, *Endozoicomonas* was typically found as a dominant microbe across a wide variety of coral species in various geographic locations. This includes *Acropora tenuis* in the Philippines, *A. millepora* in nearshore and offshore reefs in the Great Barrier Reef, and *Pocillopora verrucosa* from the Central Red Sea (Baquiran *et al*., 2025; Lema *et al*., 2014; Pogoreutz *et al*., 2018). *Endozoicomonas* relative abundance varies with thermal stress, and therefore may be a useful indicator of incipient thermal and environmental stress and should be used to monitor pre-bleaching environmental stress. Haliscomenobacteraceae, the third-best predictor of the environment, was positively associated with sea surface temperature seasonal range. This family has not been previously characterized within the context of bleaching and environmental perturbations in corals.

Together, these results suggest that shifts in the abundance of specific microbial families may serve as indicators of environmental change. This has potential utility for reef monitoring and management, and also for situations when changes in the environment cannot be measured directly (for example, in ancient DNA studies).

## 5. Future directions

We found that photosymbionts, host genetics, and the environment predict microbiome variation in central Great Barrier Reef *Acropora millepora*. The direction of influence is most certainly from environment and host genetics to microbiome, but it remains unclear whether the link between symbiont community and microbiome is similarly unidirectional (symbionts influence microbiome) or if there is a degree of feedback (microbes influence symbiont composition). The role of within-genus symbiont variation is also unclear and should be resolved with finer molecular markers. In the context of the link between microbiome and intraspecific genetic structure of the coral host, future work should aim to assess fitness consequences of microbiome shifts among diverging coral populations. We also find that the microbiome is diagnostic of environmental variation, which suggests that microbial composition can be used as a proxy of environmental change. Still, long-term tracing of coral microbiomes is needed to properly assess the diagnostic capacity of coral microbiomes to characterize incipient environmental fluctuations and heat stress, especially in threatened coral reefs such as the Great Barrier Reef and the Florida Keys.

## Acknowledgements

The data analysis for this manuscript has been performed using facilities of the Texas Advanced Computing Center (TACC). We would like to thank Zach Fuller for assistance in data curation and analytical insight. We would also like to thank Christian Voolstra for providing us with the updated photosymbiont reference genomes. This work was supported by NIH grant GM116853 to M.K.

## Data accessibility

All paired-end reads are deposited on the Sequence Read Archive under accession nos. SAMN13447100-13447355 and under BioProject PRJNA593014.

## Author contributions

- Research design: K.P., M.M.
- Performed research: K.P., M.M.
- Contributed new reagents or analytical tools: K.P., C.S., A.H., C.P., M.M.
- Analyzed data: K.P., M.M.
- Wrote the paper: K.P., M.K., M.M.
- Provided edits: A.H., C.S., C.P.
- Provided funding: K.P., M.K., M.M.

## Notes

### Competing Interest Statement

The authors have declared no competing interest.

## References

Ayre, D. J., & Hughes, T. P. (2000). Genotypic diversity and gene flow in brooding and spawning corals along the Great Barrier Reef, Australia. Evolution, 54(5), 1590–1605.

Ayre, D. J., & Hughes, T. P. (2004). Climate change, genotypic diversity and gene flow in reef building corals. Ecology Letters, 7(4), 273–278.

Baker, A. C., Glynn, P. W., & Riegl, B. (2008). Climate change and coral reef bleaching: An ecological assessment of long-term impacts, recovery trends and future outlook. Estuarine, Coastal and Shelf Science, 80(4), 435–471.

Barshis, D. J., Ladner, J. T., Oliver, T. A., Seneca, F. O., Traylor-Knowles, N., & Palumbi, S. R. (2013). Genomic basis for coral resilience to climate change. Proceedings of the National Academy of Sciences, 110(4), 1387–1392.

Barreto, M.M., Ziegler, M., Venn, A., Tambutté, E., Zoccola, D., Tambutté, S., Allemand, D., Antony, C.P., Voolstra, C.R. and Aranda, M. (2021). Effects of ocean acidification on resident and active microbial communities of Stylophora pistillata. Frontiers in Microbiology, 12, 707674.

Baquiran, J. I. P., Quijano, J. B., van Oppen, M. J., Cabaitan, P. C., Harrison, P. L., & Conaco, C. (2025). Microbiome stability is linked to Acropora coral thermotolerance in Northwestern Philippines. Environmental Microbiology, 27(2), e70041.

Beleneva, I. A., Dautova, T. I., & Zhukova, N. V. (2005). Characterization of communities of heterotrophic bacteria associated with healthy and diseased corals in Nha Trang Bay (Vietnam). Microbiology, 74(5), 579–587.

Black, K. L., Rippe, J. P., & Matz, M. V. (2025). Environmental Drivers of Genetic Divergence in Two Corals From the Florida Keys. Evolutionary Applications, 18(7), e70126.

Botté, E. S., Cantin, N. E., Mocellin, V. J., O’Brien, P. A., Rocker, M. M., Frade, P. R., & Webster, N. S. (2022). Reef location has a greater impact than coral bleaching severity on the microbiome of Pocillopora acuta. Coral Reefs, 41(1), 63–79.

Breiman, L. (2001). Random forests. Machine Learning, 45(1), 5–32.

Butler, C. C., Turnham, K. E., Lewis, A. M., Nitschke, M. R., Warner, M. E., Kemp, D. W., Hoegh Guldberg, O., Fitt, W.K., van Oppen, M.J., & LaJeunesse, T. C. (2023). Formal recognition of host generalist species of dinoflagellate (Cladocopium, Symbiodiniaceae) mutualistic with Indo Pacific reef corals. Journal of Phycology, 59(4), 698–711.

Cabacungan, G. N., Waduwara Kankanamalage, T. N., Azam, A. F., Collins, M. R., Arratia, A. R., Gutting, A. N., Matz, M.V., & Black, K. L. (2025). Cryptic coral community composition across environmental gradients. PLoS One, 20(2), e0318653.

Ceccarelli, D.M., Logan, M., Evans, R.D., Jones, G.P., Puotinen, M., Petus, C., Russ, G.R., Sinclair Taylor, T., Srinivasan, M. and Williamson, D.H. (2024). Regional scale disturbances drive long term decline of inshore coral reef fish assemblages in the Great Barrier Reef Marine Park. Global Change Biology, 30(10) e17506.

Chen, J. Z., Junker, A., Zheng, I., Gerardo, N. M., & Vega, N. M. (2024). A strong priority effect in the assembly of a specialized insect-microbe symbiosis. Applied and Environmental Microbiology, 90(10), e00818–24.

Cheng, K., Li, X., Tong, M., Jong, M.C., Cai, Z., Zheng, H., Xiao, B. and Zhou, J. (2023). Integrated metagenomic and metaproteomic analyses reveal bacterial micro-ecological mechanisms in coral bleaching. MSystems, 8(6), e00505–23.

da Silva, D. M., Costa, R., & Keller-Costa, T. (2025). A genomic view of the bacterial family Endozoicomonadaceae in marine symbioses. Communications Biology, 8(1), 1418.

De’Ath, G., Fabricius, K. E., Sweatman, H., & Puotinen, M. (2012). The 27–year decline of coral cover on the Great Barrier Reef and its causes. Proceedings of the National Academy of Sciences, 109(44), 17995–17999.

Delgadillo-Ordoñez, N., Garcias-Bonet, N., Raimundo, I., García, F. C., Villela, H., Osman, E. O., Santoro, E. P., Curdia, J., Rosado, J. G. D., Cardoso, P., Alsaggaf, A., Barno, A., Antony, C. P., Bocanegra, C., Berumen, M. L., Voolstra, C. R., Benzoni, F., Carvalho, S., & Peixoto, R. S. (2024). Probiotics reshape the coral microbiome in situ without detectable off-target effects in the surrounding environment. Communications Biology, 7(1), Article 434.

Díaz-Almeyda, E.M., Ryba, T., Ohdera, A.H., Collins, S.M., Shafer, N., Link, C., Prado-Zapata, M., Ruhnke, C., Moore, M., González Angel, A.M. and Pollock, F.J. (2022). Thermal stress has minimal effects on bacterial communities of thermotolerant Symbiodinium cultures. Frontiers in Ecology and Evolution, 10, 764086.

Dixon, G. B., Davies, S. W., Aglyamova, G. V., Meyer, E., Bay, L. K., & Matz, M. V. (2015). Genomic determinants of coral heat tolerance across latitudes. Science, 348(6242), 1460–1462.

Donner, S. D., Skirving, W. J., Little, C. M., Oppenheimer, M., & Hoegh Guldberg, O. V. E. (2005). Global assessment of coral bleaching and required rates of adaptation under climate change. Global Change Biology, 11(12), 2251–2265.

Ellis, N., Smith, S. J., & Pitcher, C. R. (2012). Gradient forests: calculating importance gradients on physical predictors. Ecology, 93(1), 156–168.

Emslie, M.J., Ceccarelli, D.M., Logan, M., Blandford, M.I., Bray, P., Campili, A., Jonker, M.J., Parker, J.G., Prenzlau, T. and Sinclair-Taylor, T.H. (2024). Changing dynamics of Great Barrier Reef hard coral cover in the Anthropocene. Coral Reefs, 43(3), 747–762.

Fuller, Z. L., Mocellin, V. J., Morris, L. A., Cantin, N., Shepherd, J., Sarre, L., Peng, J., Liao, Y., Pickrell, J., Andolfatto, P., & Matz, M. (2020). Population genetics of the coral *Acropora millepora*: Toward genomic prediction of bleaching. Science, 369(6501), eaba4674.

Gallery, D. N., Rippe, J. P., & Matz, M. V. (2025). Decrypting Corals: Does Regulatory Evolution Underlie Environmental Specialisation of Coral Cryptic Lineages?. Molecular Ecology, 34(15), e17546.

Gardner, S. G., Camp, E. F., Smith, D. J., Kahlke, T., Osman, E. O., Gendron, G., Hume, B. C. C., Pogoreutz, C., Voolstra, C. R., & Suggett, D. J. (2019). Coral microbiome diversity reflects mass coral bleaching susceptibility during the 2016 El Niño heat wave. Ecology and Evolution, 9(3), 938–956.

Glasl, B., Bourne, D. G., Frade, P. R., Thomas, T., Schaffelke, B., & Webster, N. S. (2019). Microbial indicators of environmental perturbations in coral reef ecosystems. Microbiome, 7(1), 94.

Glynn, P. W. (1996). Coral reef bleaching: facts, hypotheses and implications. Global Change Biology, 2(6), 495–509.

Grottoli, A.G., Dalcin Martins, P., Wilkins, M.J., Johnston, M.D., Warner, M.E., Cai, W.J., Melman, T.F., Hoadley, K.D., Pettay, D.T., Levas, S. and Schoepf, V. (2018). Coral physiology and microbiome dynamics under combined warming and ocean acidification. PloS One, 13(1), e0191156.

Hughes, T. P., Kerry, J. T., & Simpson, T. (2018). Large scale bleaching of corals on the Great Barrier Reef. Ecology, 99(2).

Ihaka, R., & Gentleman, R. (1996). R: a language for data analysis and graphics. Journal of Computational and Graphical Statistics, 5(3), 299–314.

Jones, A. M., Berkelmans, R., van Oppen, M. J., Mieog, J. C., & Sinclair, W. (2008). A community change in the algal endosymbionts of a scleractinian coral following a natural bleaching event: field evidence of acclimatization. Proceedings of the Royal Society B: Biological Sciences, 275(1641), 1359–1365.

Kayanne, H. (2017). Validation of degree heating weeks as a coral bleaching index in the northwestern Pacific. Coral Reefs, 36(1), 63–70.

Kellogg, C. A. (2019). Microbiomes of stony and soft deep-sea corals share rare core bacteria. Microbiome, 7(1), 90.

Korneliussen, T. S., Albrechtsen, A., & Nielsen, R. (2014). ANGSD: analysis of next generation sequencing data. BMC Bioinformatics, 15(1), 356.

Krueger, F. (2015). Trim Galore!: A wrapper around Cutadapt and FastQC to consistently apply adapter and quality trimming to FastQ files, with extra functionality for RRBS data. Babraham Institute.

Kuang, W., Li, J., Zhang, S., & Long, L. (2015). Diversity and distribution of Actinobacteria associated with reef coral Porites lutea. Frontiers in Microbiology, 6, 1094.

Lema, K. A., Bourne, D. G., & Willis, B. L. (2014). Onset and establishment of diazotrophs and other bacterial associates in the early life history stages of the coral Acropora millepora. Molecular Ecology, 23(19), 4682–4695.

Longley, R., Benucci, G. M. N., Pochon, X., Bonito, G., & Bonito, V. (2024). Species-specific coral microbiome assemblages support host bleaching resistance during an extreme marine heatwave. Science of The Total Environment, 906, 167803.

Lu, J., Breitwieser, F. P., Thielen, P., & Salzberg, S. L. (2017). Bracken: estimating species abundance in metagenomics data. PeerJ Computer Science, 3, e104.

Maire, J., Blackall, L. L., & van Oppen, M. J. (2021). Intracellular bacterial symbionts in corals: challenges and future directions. Microorganisms, 9(11), 2209.

Marangon, E., Rädecker, N., Li, J.Y., Terzin, M., Buerger, P., Webster, N.S., Bourne, D.G. and Laffy, P.W. (2025). Destabilization of mutualistic interactions shapes the early heat stress response of the coral holobiont. Microbiome, 13(1), 31.

Marshall, P. A., & Baird, A. H. (2000). Bleaching of corals on the Great Barrier Reef: differential susceptibilities among taxa. Coral Reefs, 19(2), 155–163.

Matthews, J. L., Raina, J. B., Kahlke, T., Seymour, J. R., van Oppen, M. J., & Suggett, D. J. (2020). Symbiodiniaceae bacteria interactions: rethinking metabolite exchange in reef building corals as multi partner metabolic networks. Environmental Microbiology, 22(5), 1675–1687.

Matthews, J.L., Hoch, L., Raina, J.B., Pablo, M., Hughes, D.J., Camp, E.F., Seymour, J.R., Ralph, P.J., Suggett, D.J. and Herdean, A.(2023). Symbiodiniaceae photophysiology and stress resilience is enhanced by microbial associations. Scientific Reports, 13(1), 20724.

Matz, M. V., & Black, K. L. (2024). RDAforest: identifying environmental drivers of polygenic adaptation. *Molecular Ecology Resources*, e70002.

McClanahan, T. R. (2004). The relationship between bleaching and mortality of common corals. Marine Biology, 144(6), 1239–1245.

Modolon, F., Garritano, A.D.N., Hill, L.J., Duarte, G., Bendia, A., de Moura, R.B., Pellizari, V., Thomas, T. and Peixoto, R. (2025). Promiscuous endosymbionts in deep-sea corals and crinoids are shaped by nitrogen cycling. BioRxiv, 2025-02.

Mohamed, A. R., Ochsenkühn, M. A., Kazlak, A. M., Moustafa, A., & Amin, S. A. (2023). The coral microbiome: towards an understanding of the molecular mechanisms of coral–microbiota interactions. FEMS Microbiology Reviews, 47(2), fuad005.

Morelan, I. A., Gaulke, C. A., Sharpton, T. J., Vega Thurber, R., & Denver, D. R. (2019). Microbiome variation in an intertidal sea anemone across latitudes and symbiotic states. Frontiers in Marine Science, 6, 7.

Morrow, K.M., Bourne, D.G., Humphrey, C., Botté, E.S., Laffy, P., Zaneveld, J., Uthicke, S., Fabricius, K.E. and Webster, N.S. (2015). Natural volcanic CO2 seeps reveal future trajectories for host–microbial associations in corals and sponges. The ISME Journal, 9(4), 894–908.

Naugle, M. S., Denis, H., Mocellin, V. J., Laffy, P. W., Popovic, I., Bay, L. K., & Howells, E. J. (2024). Heat tolerance varies considerably within a reef-building coral species on the Great Barrier Reef. Communications Earth & Environment, 5(1), 525.

Nishijima, M., Adachi, K., Katsuta, A., Shizuri, Y., & Yamasato, K. (2013). Endozoicomonas numazuensis sp. nov., a gammaproteobacterium isolated from marine sponges, and emended description of the genus Endozoicomonas Kurahashi and Yokota 2007. International Journal of Systematic and Evolutionary Microbiology, 63(Pt_2), 709–714.

Palladino, G., Biagi, E., Rampelli, S., Musella, M., D’Amico, F., Turroni, S., Brigidi, P., Luna, G.M. and Candela, M. (2021). Seasonal changes in microbial communities associated with the jewel anemone Corynactis viridis. Frontiers in Marine Science, 8, 627585.

Peixoto, R. S., Rosado, P. M., Leite, D. C. D. A., Rosado, A. S., & Bourne, D. G. (2017). Beneficial microorganisms for corals (BMC): proposed mechanisms for coral health and resilience. Frontiers in Microbiology, 8, 341.

Pogoreutz, C., Rädecker, N., Cárdenas, A., Gärdes, A., Wild, C., & Voolstra, C. R. (2018). Dominance of Endozoicomonas bacteria throughout coral bleaching and mortality suggests structural inflexibility of the Pocillopora verrucosa microbiome. Ecology and Evolution, 8(4), 2240–2252.

Pootakham, W., Mhuantong, W., Yoocha, T., Putchim, L., Jomchai, N., Sonthirod, C., Naktang, C., Kongkachana, W. and Tangphatsornruang, S. (2019). Heat induced shift in coral microbiome reveals several members of the Rhodobacteraceae family as indicator species for thermal stress in Porites lutea. Microbiology Open, 8(12), e935.

Primov, K. D., Burdick, D. R., Lemer, S., Forsman, Z. H., & Combosch, D. J. (2024). Genomic data reveals habitat partitioning in massive Porites on Guam, Micronesia. Scientific Reports, 14(1), 17107.

Quek, Z.R., Tanzil, J.T., Jain, S.S., Yong, W.L.O., Yu, D.C.Y., Soh, Z., Ow, Y.X., Tun, K., Huang, D. and Wainwright, B.J. (2023). Limited influence of seasonality on coral microbiomes and endosymbionts in an equatorial reef. Ecological Indicators, 146, 109878.

Rabbani, G., Afiq-Rosli, L., Lee, J. N., Waheed, Z., & Wainwright, B. J. (2025). Effects of life history strategy on the diversity and composition of the coral holobiont communities of Sabah, Malaysia. Scientific Reports, 15(1), 4459.

Richards, Z. T., & Day, J. C. (2018). Biodiversity of the Great Barrier Reef—how adequately is it protected?. PeerJ, 6, e4747.

Rippe, J. P., Dixon, G., Fuller, Z. L., Liao, Y., & Matz, M. (2021). Environmental specialization and cryptic genetic divergence in two massive coral species from the Florida Keys Reef Tract. Molecular Ecology, 30(14), 3468–3484.

Rivera, H. E., Cohen, A. L., Thompson, J. R., Baums, I. B., Fox, M. D., & Meyer-Kaiser, K. S. (2022). Palau’s warmest reefs harbor thermally tolerant corals that thrive across different habitats. Communications Biology, 5(1), 1394.

Rosado, P. M., Leite, D. C. A., Duarte, G. A. S., Chaloub, R. M., Jospin, G., Nunes da Rocha, U., Saraiva, J. P., Dini-Andreote, F., Eisen, J. A., Bourne, D. G., & Peixoto, R. S. (2019). Marine probiotics: Increasing coral resistance to bleaching through microbiome manipulation. ISME Journal, 13(4), 921–936.

Santoro, E. P., Borges, R. M., Espinoza, J. L., Freire, M., Messias, C. S., Villela, H. D., Pereira, L. M., Vilela, C. L., Rosado, J. G., Cardoso, P. M., & Rosado, P. M. (2021). Coral microbiome manipulation elicits metabolic and genetic restructuring to mitigate heat stress and evade mortality. Science Advances, 7(33), eabg3088.

Scott, C. B., Schott, R., & Matz, M. V. (2025). Genetic clustering within massive Porites species complex is the primary driver of holobiont assembly. Plos One, 20(7), e0328479.

Shan, H.W., Xia, X.J., Feng, Y.L., Wu, W., Li, H.J., Sun, Z.T., Li, J.M. and Chen, J.P.(2024). The plant-sucking insect selects assembly of the gut microbiota from environment to enhance host reproduction. Npj Biofilms and Microbiomes, 10(1), 64.

Skirving, W. J., Heron, S. F., Marsh, B. L., Liu, G., De La Cour, J. L., Geiger, E. F., & Eakin, C. M. (2019). The relentless march of mass coral bleaching: a global perspective of changing heat stress. Coral Reefs, 38(4), 547–557.

Starko, S., Fifer, J. E., Claar, D. C., Davies, S. W., Cunning, R., Baker, A. C., & Baum, J. K. (2023). Marine heatwaves threaten cryptic coral diversity and erode associations among coevolving partners. Science Advances, 9(32), eadf0954.

Stekhoven, D. J., & Bühlmann, P. (2012). MissForest—non-parametric missing value imputation for mixed-type data. Bioinformatics, 28(1), 112–118.

Sturm, A. B., Eckert, R. J., Carreiro, A. M., & Voss, J. D. (2022). Population genetic structure of the broadcast spawning coral, Montastraea cavernosa, demonstrates refugia potential of upper mesophotic populations in the Florida Keys. Coral Reefs, 41(3), 587–598.

Sturm, A. B., Eckert, R. J., Carreiro, A. M., Simões, N., & Voss, J. D. (2022). Depth-dependent genetic structuring of a depth-generalist coral and its Symbiodiniaceae algal communities at Campeche Bank, Mexico. Frontiers in Marine Science, 9, 835789.

Sturm, A. B., Eckert, R. J., Méndez, J. G., González-Díaz, P., & Voss, J. D. (2020). Population genetic structure of the great star coral, Montastraea cavernosa, across the Cuban archipelago with comparisons between microsatellite and SNP markers. Scientific Reports, 10(1), 15432.

Sully, S., Burkepile, D. E., Donovan, M. K., Hodgson, G., & Van Woesik, R. (2019). A global analysis of coral bleaching over the past two decades. Nature Communications, 10(1), 1264.

Sun, F., Yang, H., Zhang, X., Tan, F., & Shi, Q. (2022). Response characteristics of bacterial communities in multiple coral genera at the early stages of coral bleaching during El Niño. Ecological Indicators, 144, 109569.

Swain, T. D., Vega Perkins, J. B., Oestreich, W. K., Triebold, C., DuBois, E., Henss, J., Baird, A. H., Siple, M., Backman, V., & Marcelino, L. (2016). Coral Bleaching Response Index: A new tool to standardize and compare susceptibility to thermal bleaching. Global Change Biology, 22(7), 2475–2488.

Tandon, K., Lu, C.Y., Chiang, P.W., Wada, N., Yang, S.H., Chan, Y.F., Chen, P.Y., Chang, H.Y., Chiou, Y.J., Chou, M.S. and Chen, W.M. (2020). Comparative genomics: dominant coral-bacterium Endozoicomonas acroporae metabolizes dimethylsulfoniopropionate (DMSP). The ISME Journal, 14(5), pp.1290–1303.

Van Oppen, M. J., Peplow, L. M., Kininmonth, S., & Berkelmans, R. A. Y. (2011). Historical and contemporary factors shape the population genetic structure of the broadcast spawning coral, Acropora millepora, on the Great Barrier Reef. Molecular Ecology, 20(23), 4899–4914.

Van Woesik, R., Shlesinger, T., Grottoli, A. G., Toonen, R. J., Vega Thurber, R., Warner, M. E., Hulver, A.M., & Zaneveld, J. (2022). Coral bleaching responses to climate change across biological scales. Global Change Biology, 28(14), 4229–4250.

Voolstra, C. R., Raina, J. B., Dörr, M., Cárdenas, A., Pogoreutz, C., Silveira, C. B., Mohamed, A. R., Bourne, D. G., Luo, H., Amin, S. A., & Peixoto, R. S. (2024). The coral microbiome in sickness, in health and in a changing world. Nature Reviews Microbiology, 22(8), 460–475.

Wang, X., Feng, H., Wang, Y., Wang, M., Xie, X., Chang, H., Wang, L., Qu, J., Sun, K., He, W. and Wang, C.(2021). Mycorrhizal symbiosis modulates the rhizosphere microbiota to promote rhizobia–legume symbiosis. Molecular Plant, 14(3), 503–516.

Wedin, M., Maier, S., Fernandez Brime, S., Cronholm, B., Westberg, M., & Grube, M. (2016). Microbiome change by symbiotic invasion in lichens. Environmental Microbiology, 18(5), 1428–1439.

West, J. M., & Salm, R. V. (2003). Resistance and resilience to coral bleaching: implications for coral reef conservation and management. Conservation Biology, 17(4), 956–967.

Wood, D. E., Lu, J., & Langmead, B. (2019). Improved metagenomic analysis with Kraken 2. Genome Biology, 20(1), 257.

Xu, M., Cheng, K., Xiao, B., Tong, M., Cai, Z., Jong, M.C., Chen, G. and Zhou, J. (2023). Bacterial communities vary from different scleractinian coral species and between bleached and non-bleached corals. Microbiology Spectrum, 11(3), e04910–22.

Xu, Y., Liang, J., Qin, L., Niu, T., Liang, Z., Li, Z., Chen, B., Zhou, J. and Yu, K. (2024). The dynamics of Symbiodiniaceae and photosynthetic bacteria under high-temperature conditions. Microbial Ecology, 87(1), 169.

Yang, Q., Zhang, H., Qiu, J. W., Huang, D., Zhou, X., & Zheng, X. (2025). Symbiotic Symbiodiniaceae mediate coral-associated bacterial communities along a natural thermal gradient. Environmental Microbiome, 20(1), 72.

Ziegler, M., Grupstra, C.G., Barreto, M.M., Eaton, M., BaOmar, J., Zubier, K., Al-Sofyani, A., Turki, A.J., Ormond, R. and Voolstra, C.R. (2019). Coral bacterial community structure responds to environmental change in a host-specific manner. Nature Communications, 10(1), 3092.

