## Supplementary Tables and Figures for "*Acropora millepora*’s microbiome is predicted by algal symbionts, host genetics, and environment"

MOLECULAR ECOLOGY

### Table of Contents:


| | |
|---|---|
| **Figure S1: Environmental variable clustering dendrogram** | Page 2 |
| **Figure S2: Microbial family clustering dendrogram** | Page 3 |
| **Figure S3: Top 15 most abundant microbial families** | Page 4 |
| **Figure S4: Microbiome ordination** | Page 5 |
| **Figure S5: Microbiome RDAforest** | Page 6 |
| **Figure S6: Microbiome turnover curves** | Page 7 |
| **Figure S7: RDAforest microbiome model performance drop** | Page 8 |
| **Figure S8: Host genetic RDAforest** | Page 9 |
| **Figure S9: Host genetic ordination** | Page 10 |
| **Figure S10: *Endozoicomonas* RDAforest** | Page 11 |
| **Figure S11: *Endozoicomonas* turnover curves** | Page 12 |
| **Table S1: Pearson and Spearman-rho correlations** | Page 13 |
| **Table S2: RDAforest Model Importances** | Page 14 |
| **Table S3: Non-parametric regression** | Page 14 |

MOLECULAR ECOLOGY

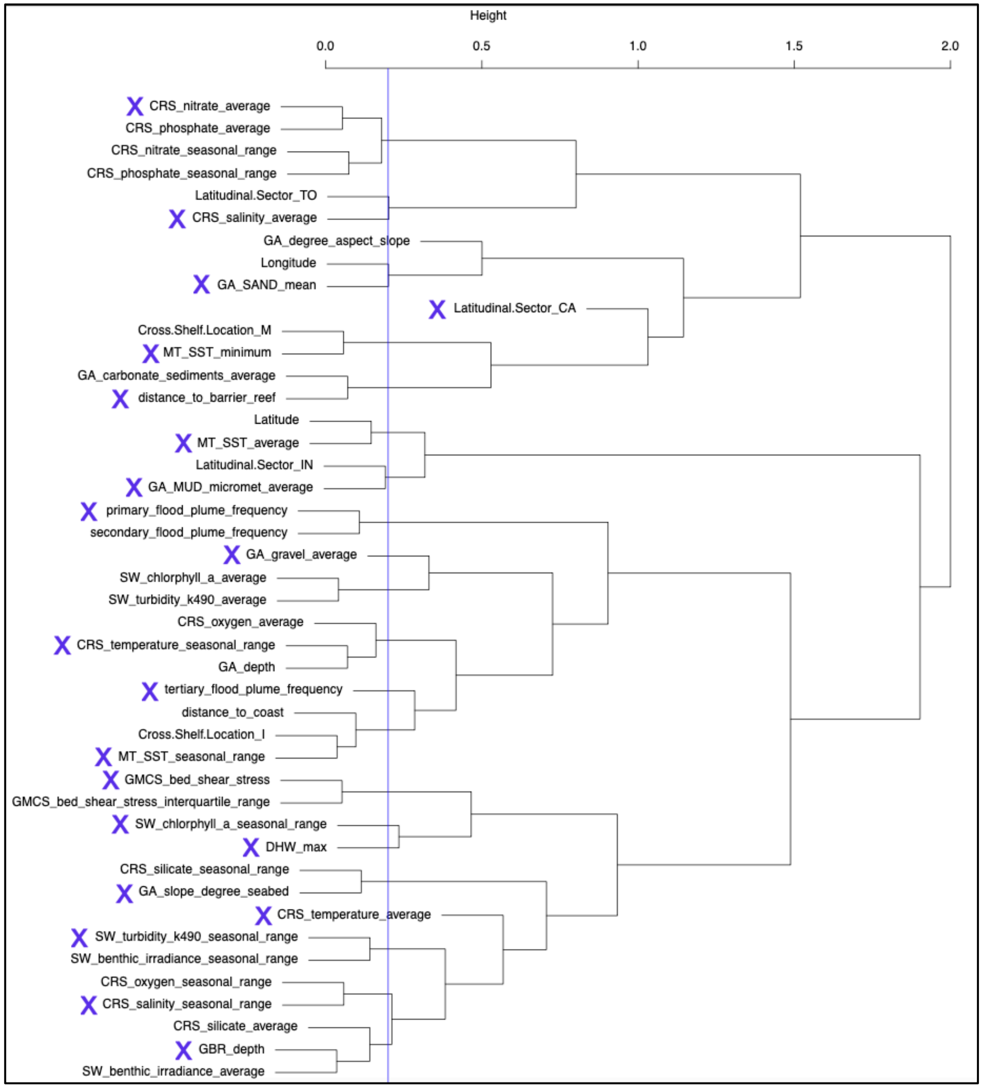


**Figure S1:** Selection of environmental variables (X-marked) that have a correlation coefficient of 0.8 or less to one another.

MOLECULAR ECOLOGY

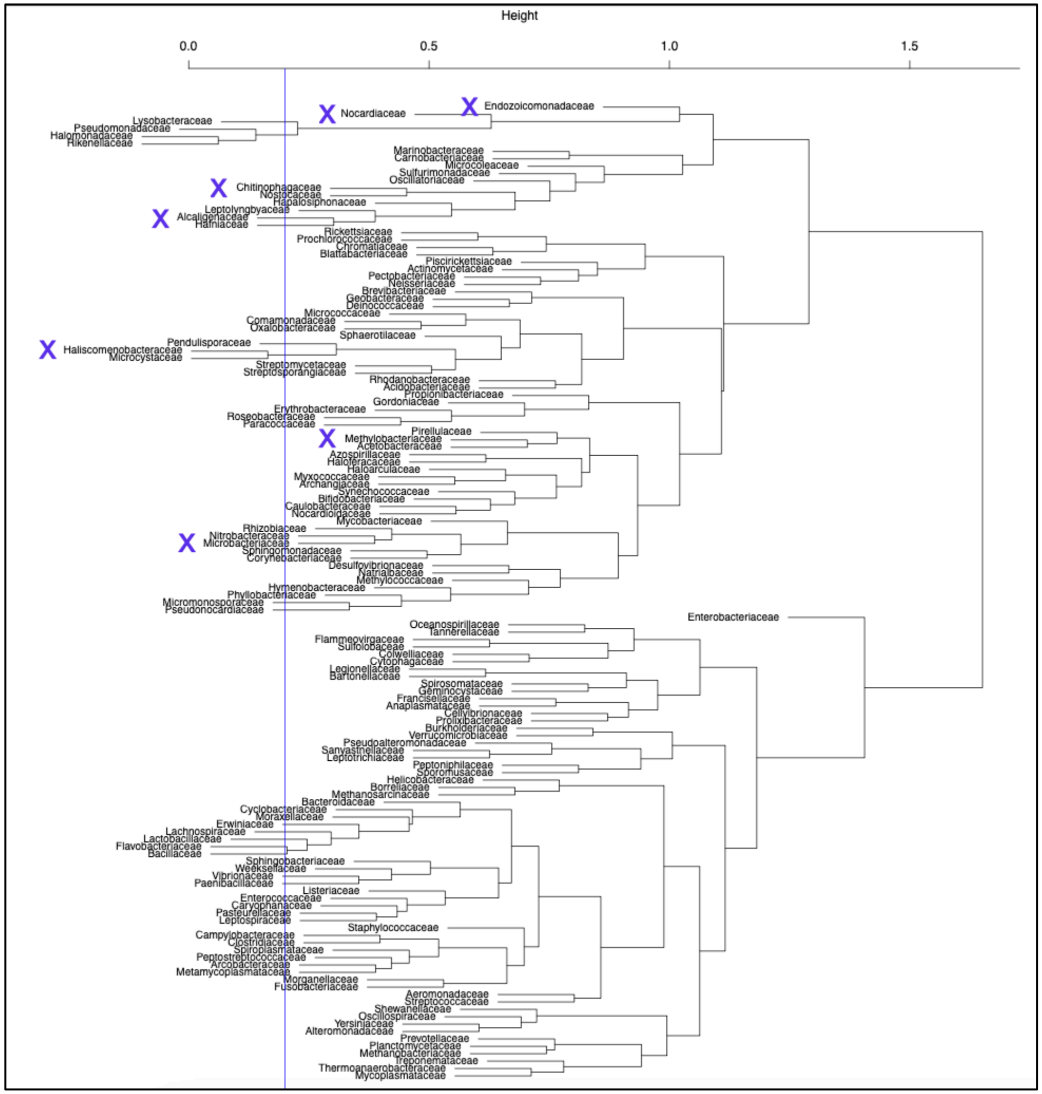


**Figure S2**: Selected microbial predictors of environmental variation (X-marked) have less than 0.8 correlation to one another.

MOLECULAR ECOLOGY

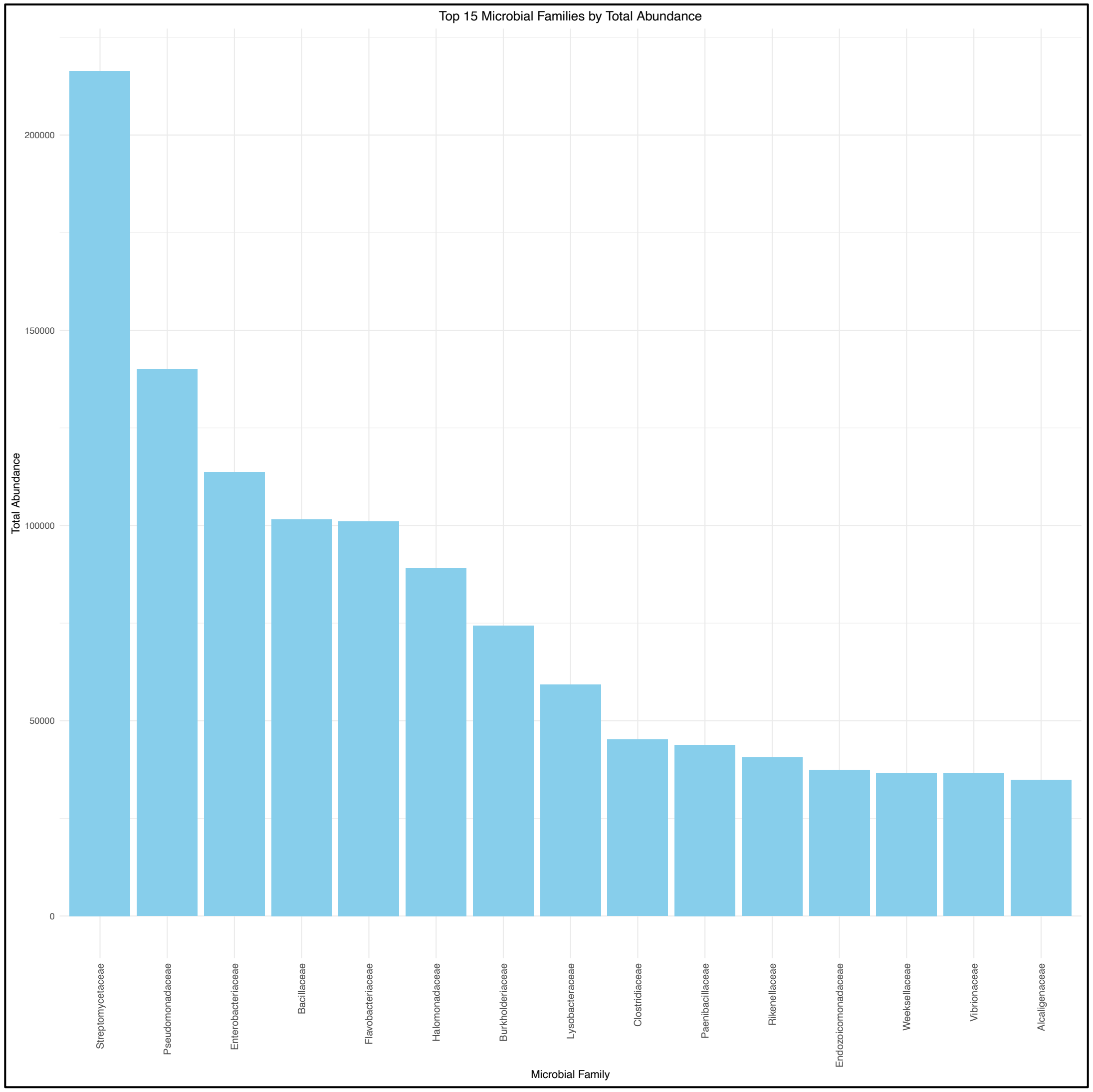


**<u>Figure S3:</u>** Top 15 most abundant microbial families of *Acropora millepora*.

MOLECULAR ECOLOGY

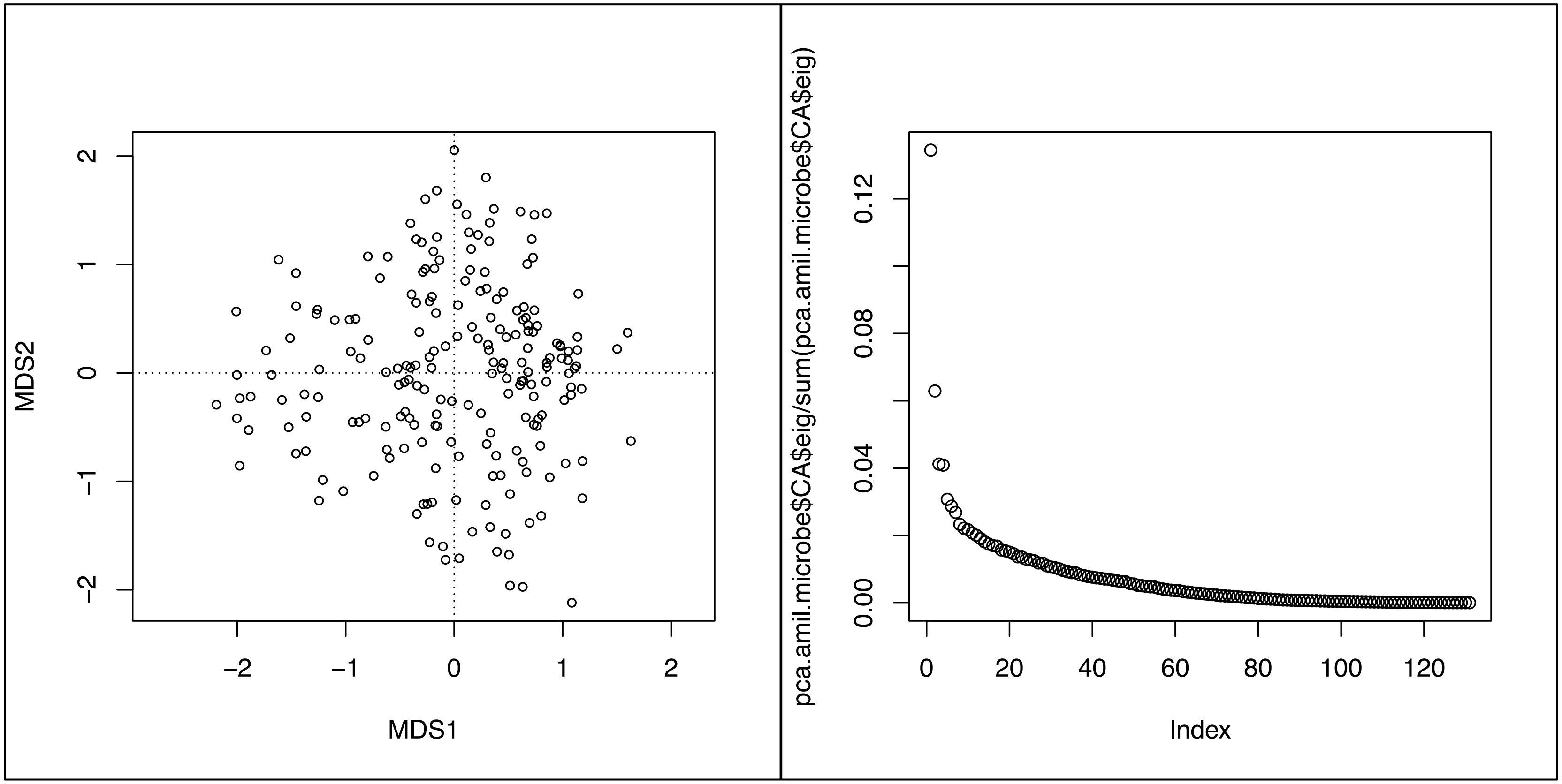


**<u>Figure S4</u>**: *Acropora millepora* microbiome unconstrained ordination (left) and screeplot (right).

MOLECULAR ECOLOGY

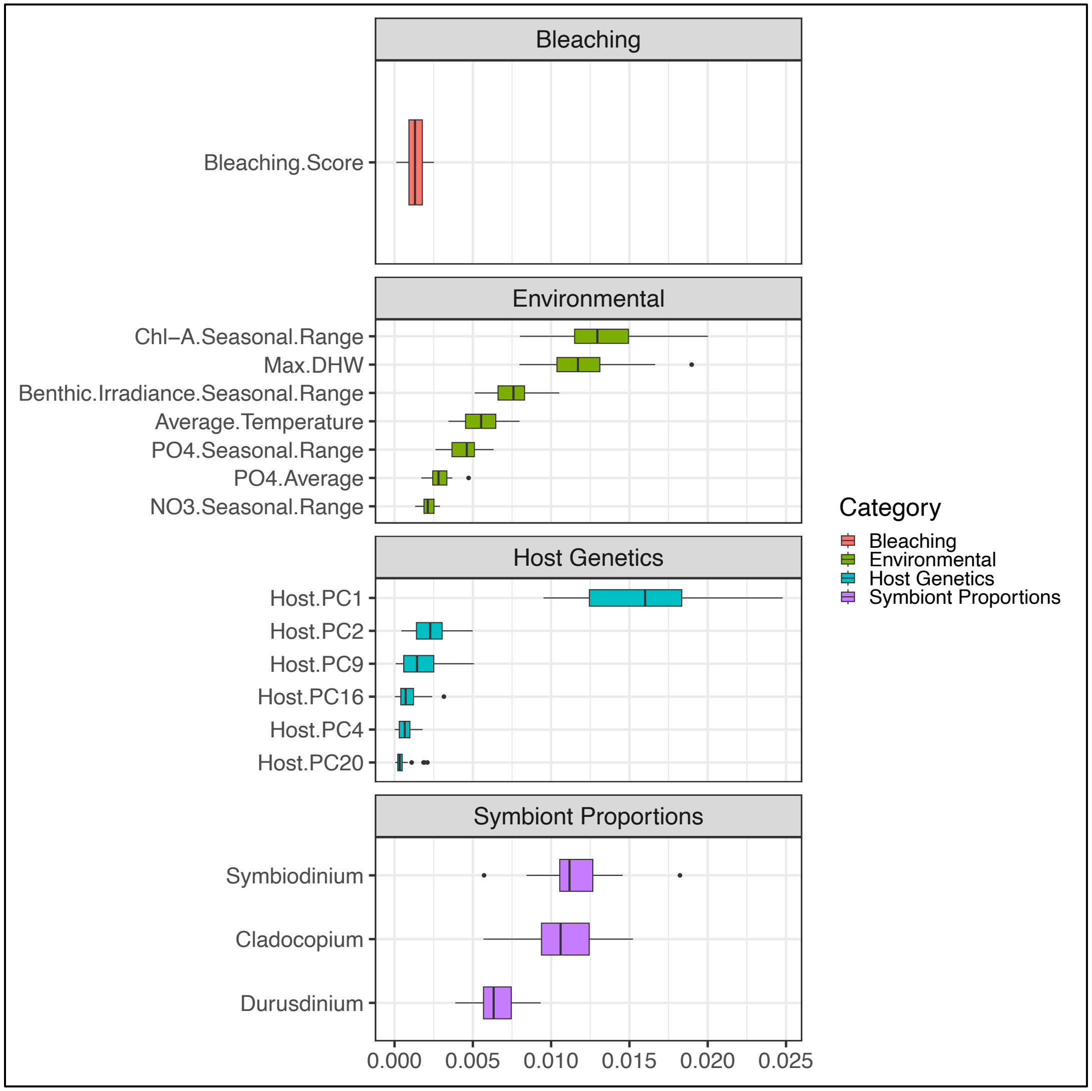


**<u>Figure S5</u>:** RDAForest ordination jackknifing confidence intervals of the top predictors of *Acropora millepora* microbiome variation.

MOLECULAR ECOLOGY

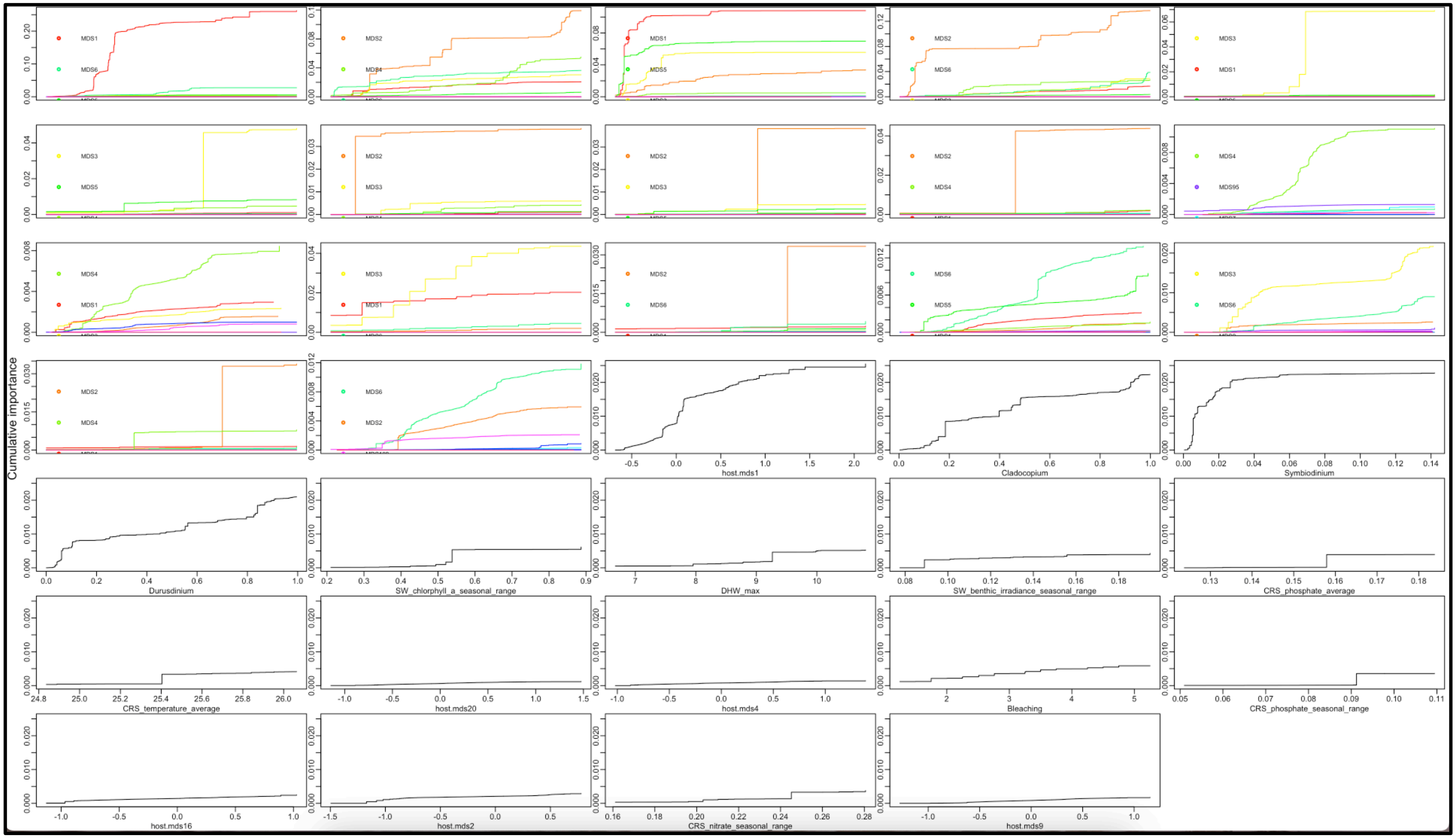


**<u>Figure S6</u>**: *Acropora millepora* microbiome turnover curves, revealing where changes in the microbiome occur at specific values for the top predictors of microbiome variation.

MOLECULAR ECOLOGY

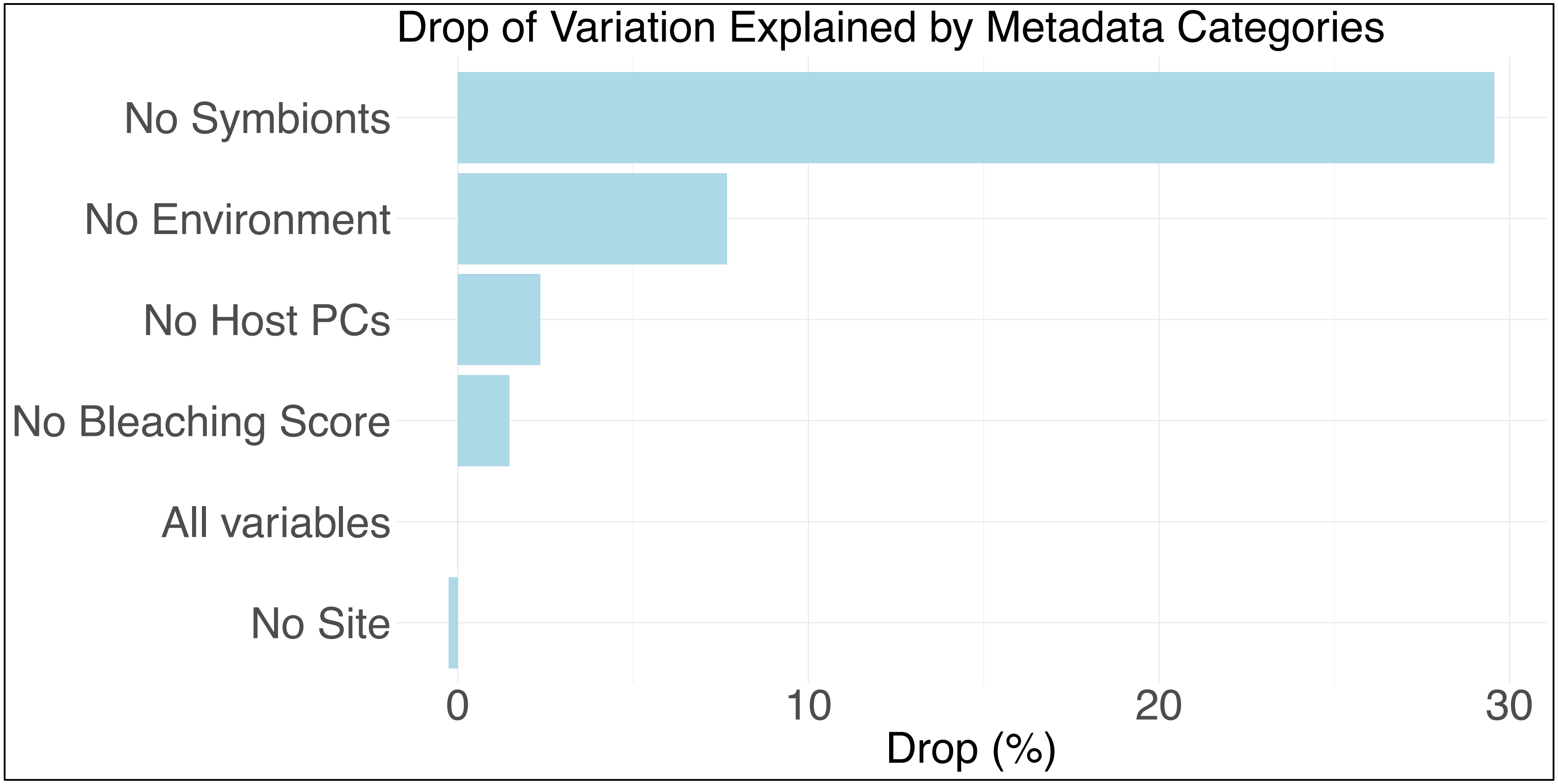


**Figure S7:** Percent drop in model variation captured by RDAForest's *makeGF* gradientForest function when excluding different metadata categories used to explain microbiome variation. Exclusion of symbiont proportions leads to the largest drop in microbiome variation captured by the gradientForest function.

MOLECULAR ECOLOGY

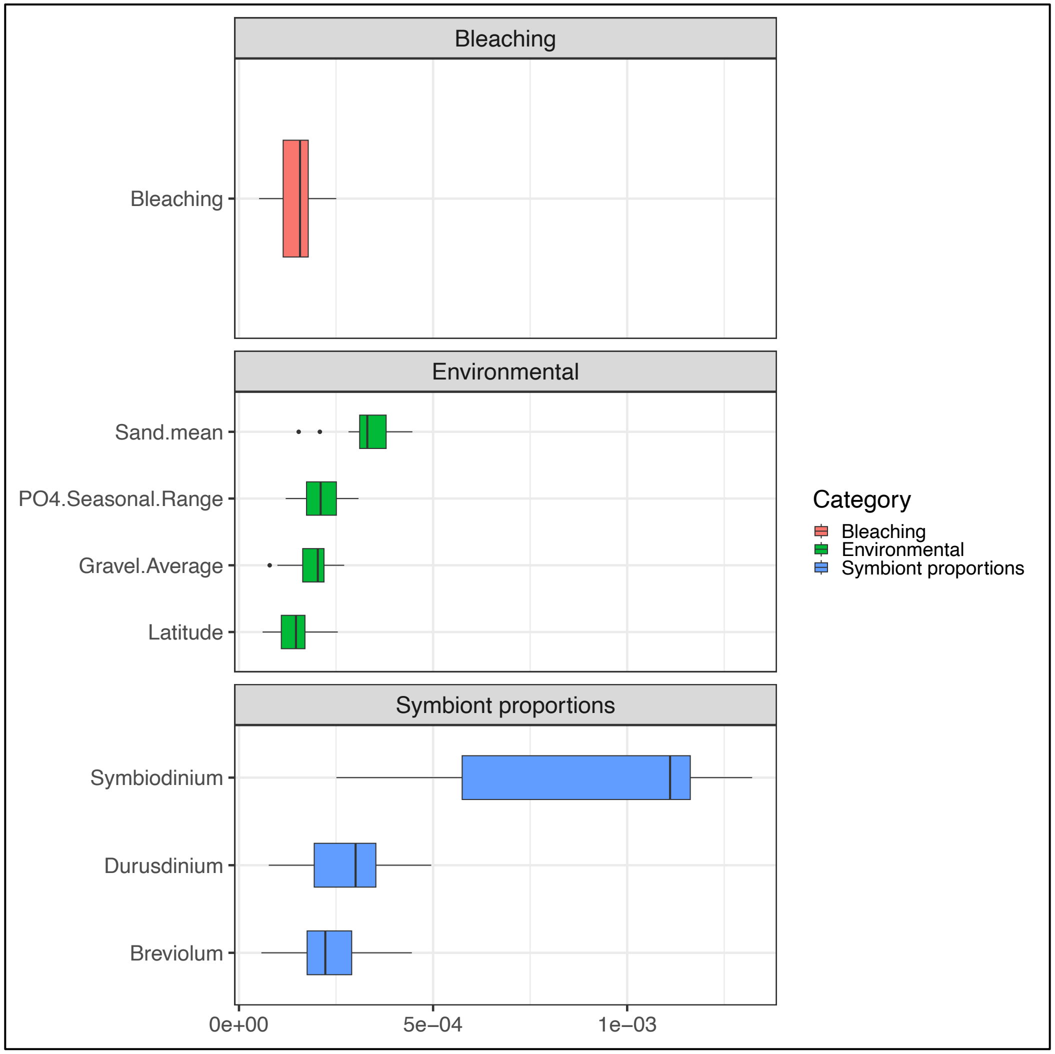


**Figure S8:** RDAForest ordination jackknifing confidence intervals of the top predictors of *Acropora millepora* host genetic variation.

MOLECULAR ECOLOGY

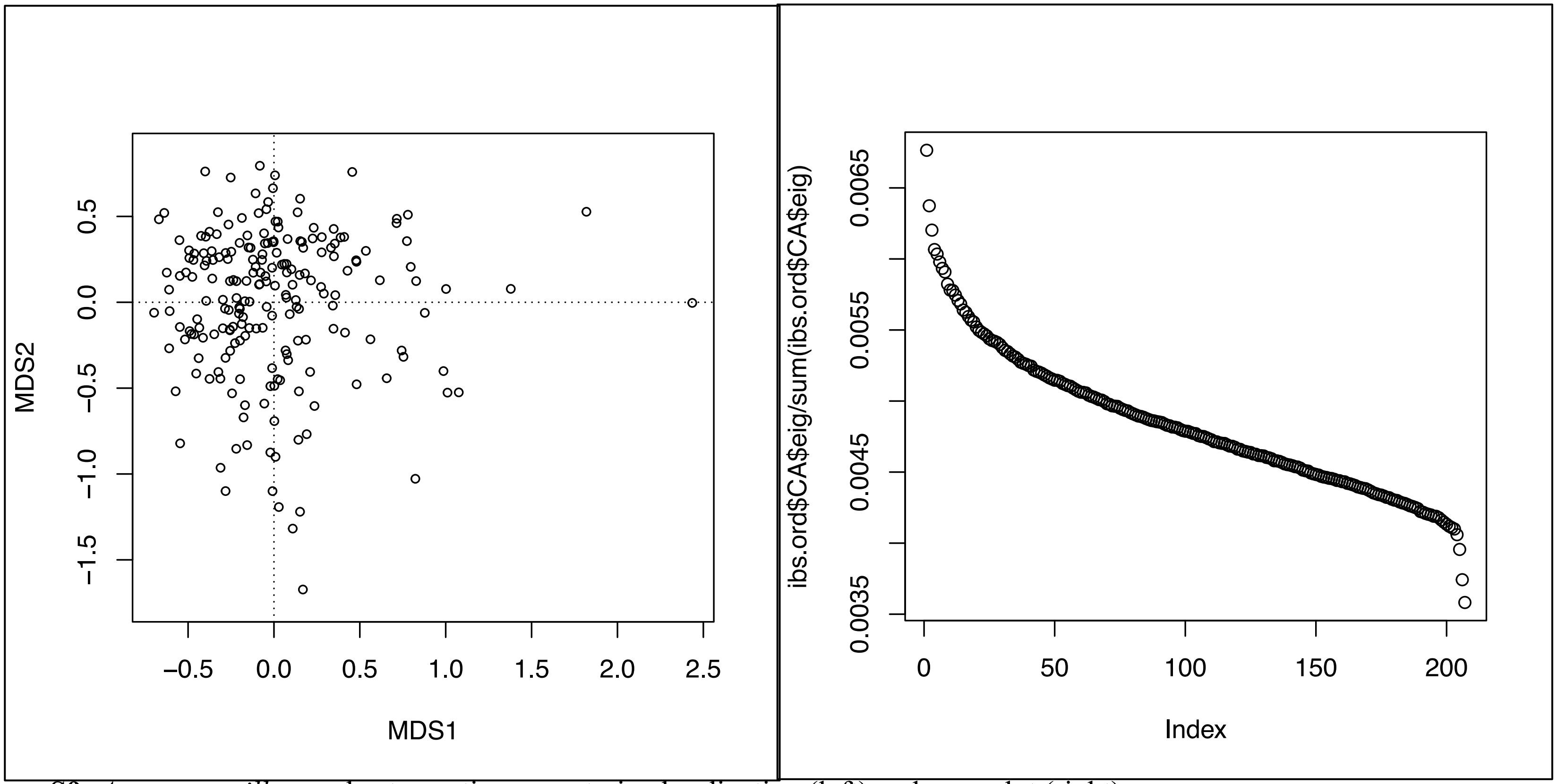


**<u>Figure S9</u>**: *Acropora millepora* host genetic unconstrained ordination (left) and screeplot (right).

MOLECULAR ECOLOGY

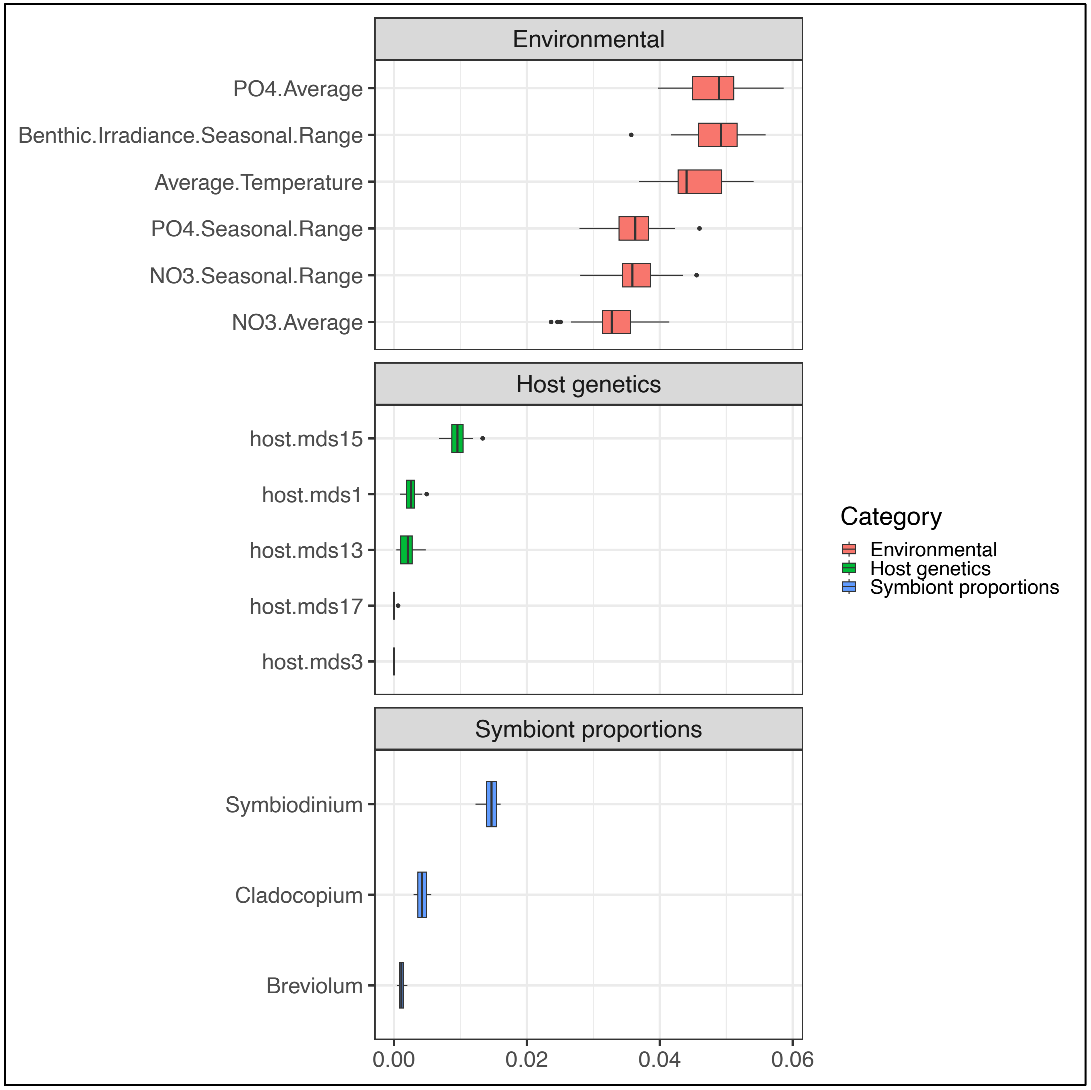


**Figure S10**: RDAForest ordination jackknifing confidence intervals of the top predictors of *Endozoicomonas* variation.

MOLECULAR ECOLOGY

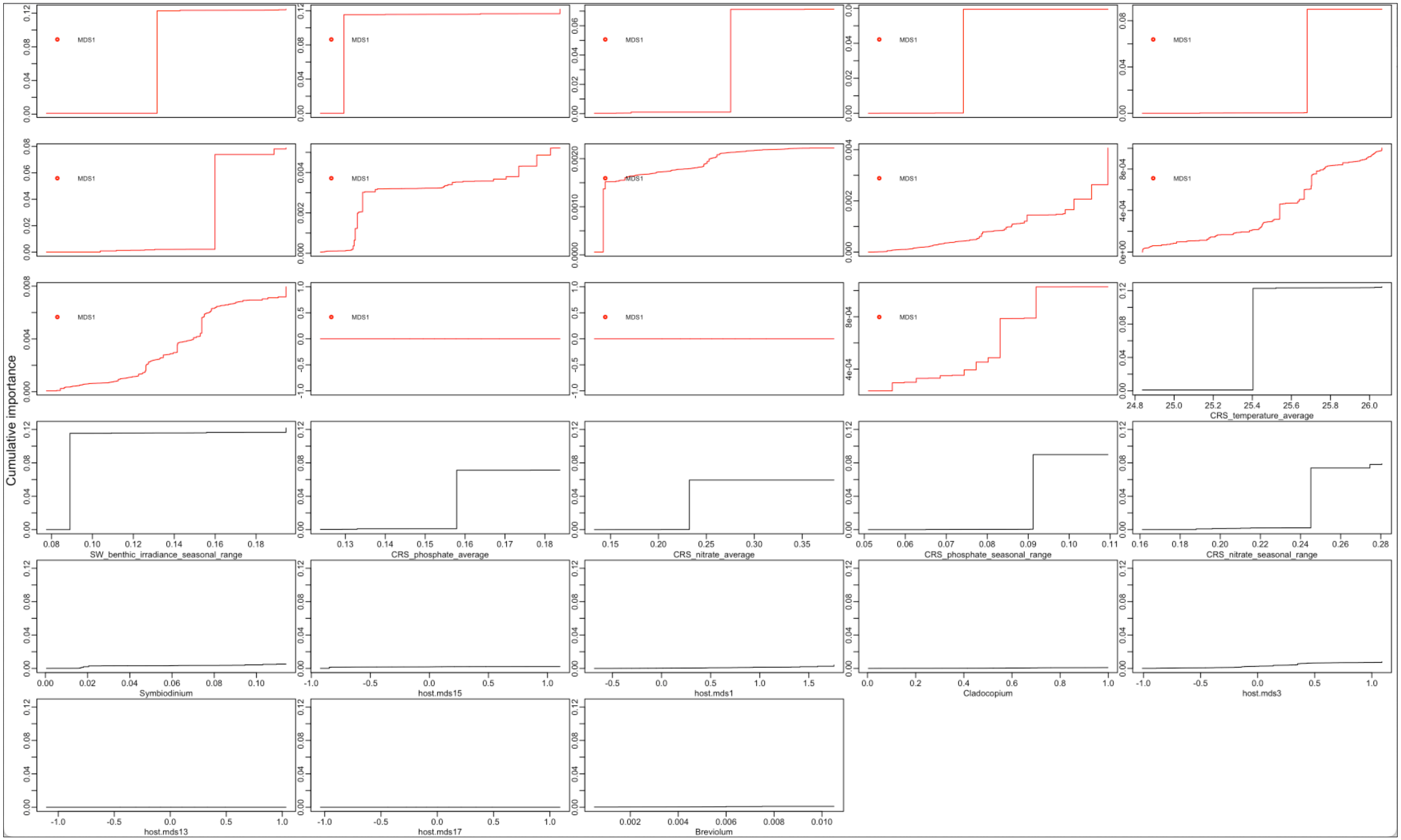


**Figure S11**: *Endozoicomonas* turnover curves, revealing where changes in *Endozoicomonas* variation occur at specific values for the top predictors of *Endozoicomonas* variation.

MOLECULAR ECOLOGY

**Table S1:** Pearson and Spearman-rho correlations of bleaching scores, *Endozoicomonas* variation, and host genetic principal component 1 to symbiont proportions, environmental variables and microbiome variation. The T value indicates a positive or negative correlation, while the p-value indicates the statistical significance of the correlation.

| | | **Pearson** | | | | **Spearman rho** | | | |
|---|---|---|---|---|---|---|---|---|---|
| **Variables Included In Correlation** | | **T** | **DF** | **p-value** | **Correlation** | **S-value** | **Rho** | **p-value** | **N** |
| **Bleaching** | Symbiodinium | -2.5334 | 206 | 0.01204 | -0.1738252 | 903115 | -0.2063087 | 0.007847 | 208 |
| | Breviolum | -3.1057 | 206 | 0.002166 | -0.2114888 | 919202 | -0.2277962 | 0.003254 | 208 |
| | Cladocopium | -4.5916 | 206 | 7.65E-06 | -0.3046989 | 713124 | 0.0474657 | 0.5449 | 208 |
| | Durusdinium | 4.7778 | 206 | 3.36E-06 | 0.3158474 | 777648 | -0.0387198 | 0.6215 | 208 |
| | Degree Heating Weeks | -5.0512 | 206 | 9.64E-07 | -0.3319742 | 1044342 | -0.3949485 | 1.52E-07 | 208 |
| **Endozoicomonas** | Degree Heating Weeks | - | - | - | - | 1843572 | -0.2292252 | 0.000867 | 208 |
| | Bleaching | - | - | - | - | 555711 | 0.2577265 | 0.0008317 | 208 |
| | Durusdinium | - | - | - | - | 1901718 | -0.2679943 | 9.09E-05 | 208 |
| | Average Temperature | - | - | - | - | 2050913 | -0.3674723 | 4.76E-08 | 208 |
| **Host genetic PC1** | Symbiodinium | -2.4067 | 206 | 0.01698 | -0.1653732 | 1845656 | -0.2306148 | 0.0008046 | 208 |
| | Breviolum | -2.9511 | 206 | 0.003533 | -0.2014006 | 1906049 | -0.2708823 | 7.58E-05 | 208 |
| | Cladocopium | 4.031 | 206 | 7.82E-05 | 0.2703945 | 972399 | 0.3516405 | 1.91E-07 | 208 |
| | Durusdinium | -3.8825 | 206 | 0.0001392 | -0.261124 | 2071499 | -0.3811985 | 1.34E-08 | 208 |
| | Microbiome PC1 | 11.137 | 206 | 2.20E-16 | 0.6130466 | 478178 | 0.6811688 | 2.20E-16 | 208 |

MOLECULAR ECOLOGY

**<u>Table S2</u>:** Drop in the R2 importances of RDAForest's *makeGF* gradientForest model when excluding different metadata categories. Exclusion of symbiont proportions leads to the largest drop in microbiome variation captured by the gradient forest function.

| Metadata Group | Full Model $R^2$ | $R^2$ (Excluded) | $R^2$ Drop Rank |
|---|---|---|---|
| **Symbionts** | 0.1568091 | 0.1104439 | 1 |
| **Environment** | 0.1568091 | 0.1447774 | 2 |
| **Host Genetics** | 0.1568091 | 0.1531245 | 3 |
| **Bleaching** | 0.1568091 | 0.154493 | 4 |
| **Population** | 0.1568091 | 0.1572145 | 5 |

**<u>Table S3</u>:** Non-parametric regressions of bleaching scores and *Endozoicomonas* variation to symbiont proportions and environmental variables. EDF = effective degrees of freedom; REF.DF = reference degrees of freedom.

| Variables Included in Regression | | EDF | REF.DF | F | p-value | N |
|---|---|---|---|---|---|---|
| **Bleaching** | Symbiodinium | 1.748 | 2.198 | 4.15 | 0.0142 | 208 |
| | Breviolum | 1.793 | 2.258 | 5.02 | 0.00523 | 208 |
| | Cladocopium | 8.306 | 8.861 | 5.453 | 2.21E-06 | 208 |
| | Durusdinium | 8.319 | 8.866 | 5.647 | 1.60E-06 | 208 |
| | Degree Heating Weeks | 7.136 | 7.993 | 4.961 | 1.62E-05 | 208 |
| **Endozoicomonas** | Degree Heating Weeks | 8.667 | 8.944 | 46.63 | <2E-16 | 208 |
| | Bleaching | 7.303 | 8.296 | 5.273 | 6.29E-06 | 208 |
| | Durusdinium | 3.271 | 3.98 | 3.702 | 0.00576 | 208 |
| | Average Temperature | 6.658 | 7.255 | 57.17 | <2E-16 | 208 |
